# Master regulators governing protein abundance across ten human cancer types

**DOI:** 10.1101/2024.11.11.619147

**Authors:** Zishan Wang, Megan Wojciechowicz, Jordan Rosen, Abdulkadir Elmas, Won-Min Song, Yansheng Liu, Kuan-lin Huang

## Abstract

Protein abundance correlates only moderately with mRNA levels, and are modulated post-transcriptionally by a network of regulators including ribosomes, RNA-binding proteins (RBPs), and the proteasome. Here, we identified **Ma**ster **P**rotein abundance **R**egulators (MaPRs) across ten cancer types by devising a new computational pipeline that jointly analyzed transcriptomes and proteomes from 1,305 tumor samples. We identified 232 to 1,394 MaPRs per cancer type, mediating up to 79% of post-transcriptional regulatory networks. MaPRs exhibit high network connectivity, strong genetic dependency in cancer cells, and significant enrichment for RBPs. Combining tumor up-regulation, druggability, and target network analyses identified cancer-specific vulnerabilities. MaPRs predict tumor proteomic subtypes more accurately than other proteins. Finally, significant portions of RBP MaPR-target relationships were validated by experimental evidence from eCLIP binding and knockdown assays. Our findings uncover central MaPRs that govern post-transcriptional networks, highlighting diverse processes underlying human proteome regulation and identifying key regulators in cancer biology.

## INTRODUCTION

Proteins execute most cellular functions. Dysregulation of proteins, such as overexpression of oncogenic kinases, diminished expression of DNA damage repair proteins, and imbalanced metabolic enzyme levels, can drive and sustain tumorigenesis. A myriad of post transcriptional processes that modulate translation rates, protein transport, or protein degradation are known to impact protein abundance^1,2^. Despite extensive research on the regulation of mRNA levels, the control of protein abundance and such key regulators in large-scale cancer cohorts remain sparsely characterized. Moreover, most cancer drug targets are proteins^3,4^, highlighting the urgent need for a systematic network approach that can leverage the emerging proteogenomic datasets to characterize post-transcriptional regulatory networks and identify key regulators of protein abundance that shape individual tumor proteomes.

Recent advancement in high-throughput mass spectrometry (MS) technologies have enabled the quantification of over ten thousand proteins from primary tumor samples^5–15^. Several of the cancer protegenomic studies that concurrently conducted global proteomics and transcriptomic analyses have reported overall moderate correlations between mRNA levels and protein abundance, with median correlations ranging from 0.39 to 0.54^16–18^ and significant variations across different genes. Discordance between mRNA and protein levels have also been reported across diverse physiological conditions and organisms^1^. These observations, coupled with the successful expansion of cancer therapeutic targets through proteogenomics analysis^3^ and identification of potential transcriptional master regulators as cancer therapeutic targets^19,20^, underscore the necessity for systematic and in-depth investigations into master regulators of protein abundance across cancer types.

We hypothesize that there exists Master Protein abundance Regulators (MaPRs) that orchestrate post-transcriptional processes regulating abundance of target proteins and shape the molecular phenotypes of cancer. Leveraging 10 cancer cohorts with spontaneous quantitative transcriptomic and proteomic profiling, we developed a computational pipeline to systematically identify MaPRs affecting protein abundance in post-transcriptional regulatory networks through diverse cellular processes. Our analysis revealed that MaPRs show high degree of network connectivity and are significantly enriched for known translational processes including RBPs and the nuclear core complex. We identified MaPR pairs with shared targets and present as cancer-specific vulnerabilities. MaPR proteins performs superiorly at predicting proteomic subtype across cancer types compared to non-MaPRs, suggesting their central role in shaping tumor proteomes. Finally, we validated a significant fraction of MaPR-target relationships using experimental RNA binding and knockdown data. Overall, our MaPR pipeline provides a new method to identify key regulators of protein abundance, illuminating their pivotal roles of determining the cancer proteome landscape.

## RESULTS

### Systematic identification of MaPRs across ten cancer types

We devised a computational pipeline, MaPR, to systematically predict master regulators of protein abundance across 10 cancer types using datasets compiled from the Clinical Proteomic Tumor Analysis Consortium (CPTAC) phase 3 studies. The curated proteogenomic dataset includes a total of 1,047 samples in the discovery set and 258 samples in the validation set (**Fig. 1A and Table S1, Methods**). The MaPR pipeline consists of four steps (**Methods**), built into an R package (**Data and code availability, Fig. 1B**). (i) mRNA and protein co-expression networks are separately constructed based on Spearman correlation. (ii) To eliminate transcriptional influence, a post-transcriptional regulatory network is generated by directly removing the edges of mRNA co-expression network from the protein co-expression network. (iii) MaPR builds a directed post-transcriptional regulatory network, where edges originate from regulators to targets. This is achieved by calculating semi-partial Spearman correlation^21^, which identifies regulators whose protein abundance correlates significantly with target protein abundance independent of target mRNA levels. Significant edges are retained to obtain the final post-transcriptional regulatory network for each cancer cohort. Across cancers, 5.3%-24.1% of significant edges are supported by the protein physical interaction network from string^22^ (**Fig. 1C and Table S1)**. Examination of the target number of all regulators revealed a power law distribution, indicating the post-transcriptional regulatory network is scale-free-like, similar to most biological networks (**Fig. S1A**). (iv) The post-transcriptional regulatory network is randomly shuffled 10,000 times to identify MaPRs that show significant regulations in the network. To ensure robustness, the network shuffling process is repeated 100 times and the overlapping MaPRs across all repetitions are defined as the final set of MaPRs (**Fig. 1B and S1B**). As an illustrative example of a MaPR-target pair, the protein abundance of the MaPR AQR showed a significant high correlation with target PAD21 protein abundance independent of *PAD21* mRNA expression in the GBM cohort (semi-partial spearman correlation = 0.725, BH-corrected P-value < 1e-6), despite of the lack of correlation between AQR protein and *PAD21* mRNA levels (**Fig. 1D**). The comprehensive list of MaPRs across cancer types is provided as a resource at **Table S1**.

**Figure 1.**
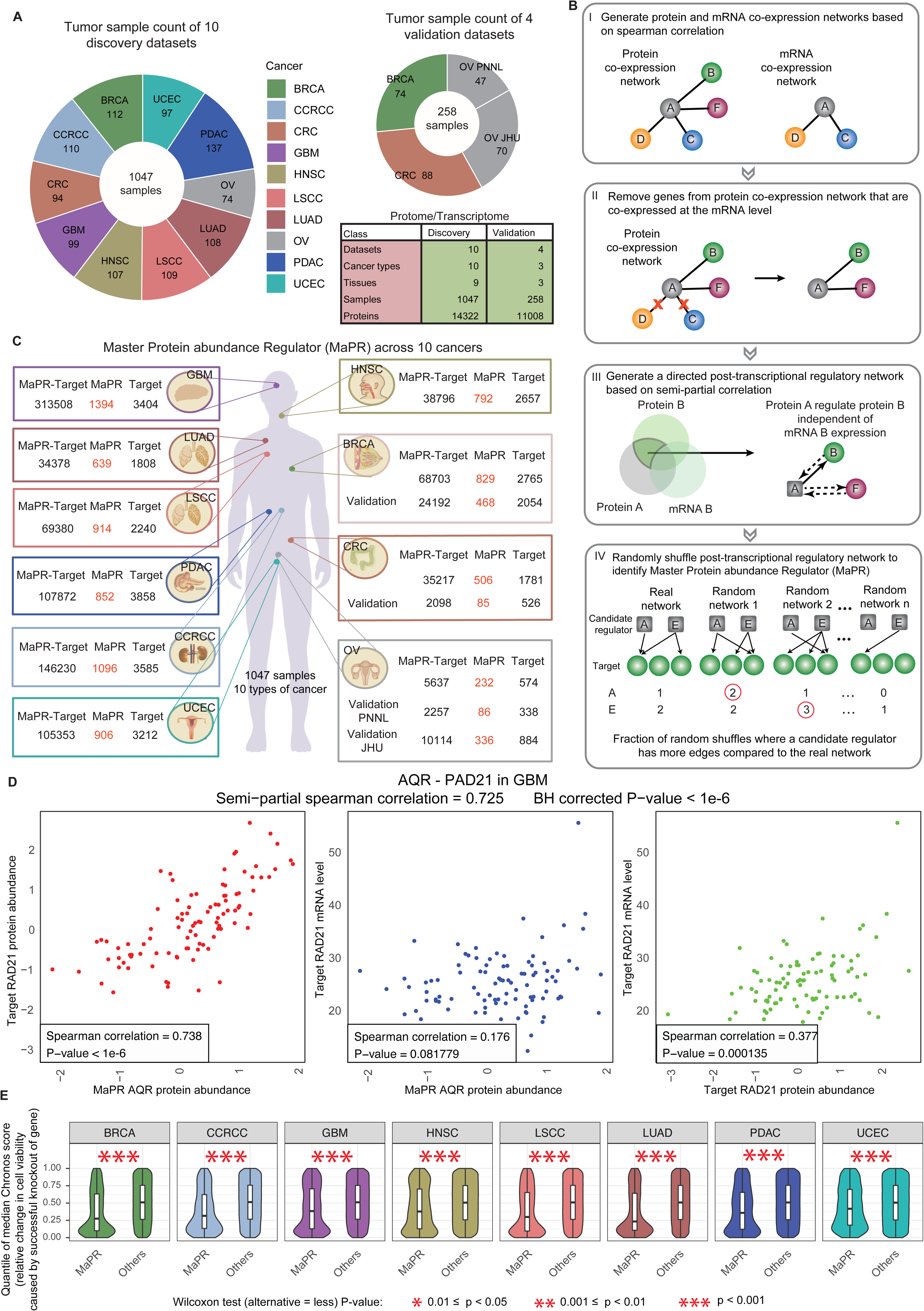
A computationally integrative pipeline to predict Master Protein abundance Regulator (MaPR) in cancer. (A) Tumor sample count with paired mRNA and protein expression profile of 14 datasets, including 10 discovery datasets of 10 cancer types and 4 validation datasets of 3 cancer types. Different colors represent different cancer types. (B) Schematic overview of the pipeline. (C) Number of identified post-transcriptional regulations, MaPR, and their targets across 10 cancers. (D) Demonstration of one MaPR-target pair through correlations between MaPR AQR protein abundance, target PAD21 mRNA level, and target PAD21 protein abundance in GBM. (E) Comparison between the quantile of median Chronos score (relative change in cell viability caused by successful CRISPR knockout) of MaPRs and other genes in corresponding cancer lines from DepMap 24Q2. A lower Chronos score indicates a loss of cancer cell viability upon knockout and higher genetic dependency. Only cancer types with corresponding cancer lines available from DepMap 24Q2 are shown. P-value was calculated by one-sided Wilcoxon test for paired median Chronos score groups and significance was denoted by asterisks in each cancer.

In total, we identified 232 to 1394 MaPRs in each of the 10 cancer types (**Fig. 1C**). While MaPRs identified by step (iv) consituted only a small proportion of the regulators of (iii), varing from 13% in OV to 25% in GBM, MaPRs of (iv) typically mediated the majority of the post-transcriptional regulatory networks of (iii)—ranging from 48% in OV to 79% in GBM (**Fig. S1C**). MaPRs was observed to regulate significantly more targets and exhibit higher Pagerank centrality score compared with other non-MaPR regulators (Wilcoxon test P-value < 0.001, **Fig. S1D and S1E**). Moreover, CRISPR genetic knockout of MaPRs reduced cell viability at larger extents compared with knocking out other genes across cancer types based on the DepMap 24Q2 dataset^23^, demonstrating MaPRs’ essentiality in cancer cells (Wilcoxon test P-value < 0.001, **Fig. 1E and S1F, Method**). To validate our predictions, the MaPR pipeline was applied to 4 independent validation datasets from CPTAC2/TCGA retrospective studies across 3 cancer types, including BRCA, CRC, and OV (OV analyzed at Pacific Northwest National Laboratory (PNNL) and Johns Hopkins University (JHU)) (**Fig. 1A and Table S1**). Subsequently, we computed cross-cancer Jaccard/geometric indices based on overlapping MaPRs (**Fig. 2A and S2C**) and edges in the post-transcriptional networks (**Fig. S2A and S2B**). As expected, post-transcriptional regulations and MaPRs from validation studies exhibited higher overlaps within the same cancer type from the CPTAC3 cohorts (**Fig. 2A**). Cancers originating from similar tissues are known to possess common molecular characteristics that can lead to tissue-specific context of protein abundance control^24–26^. Here, LUAD and LSCC, two subtypes of non-small cell lung cancer, also displayed higher overlaps in MaPRs and associated networks (**Fig. 2A and S2**), further validating that the MaPR pipeline accurately recapitulates the underlying post-transcriptional biology within tissue types.

**Figure 2.**
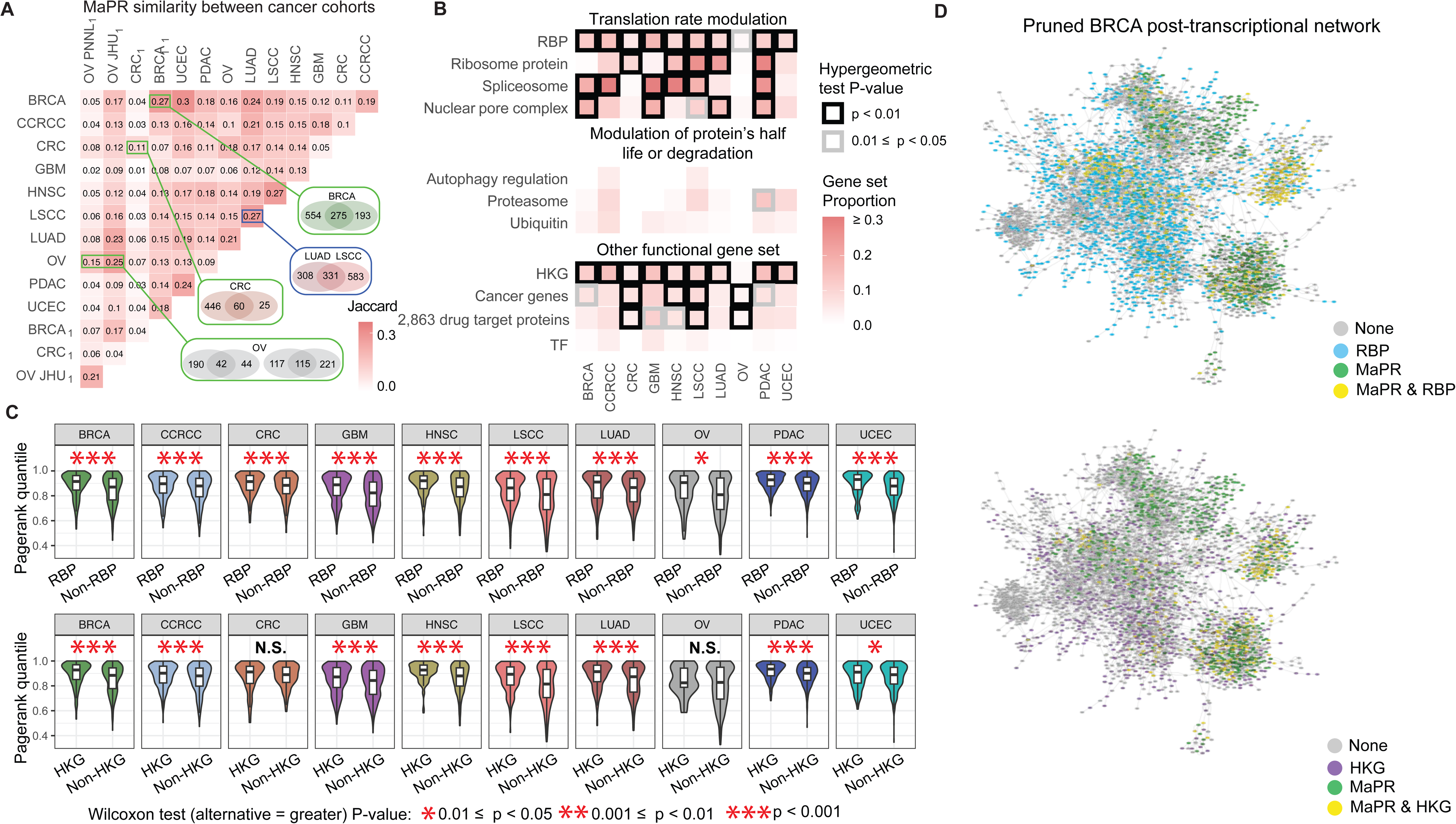
Identification of MaPR across 10 cancer types. (A) The Jaccard index matrix showed the similarity of MaPRs found across cancer types. The same cancer type from independent cohorts was labeled by green box while cancer from the same tissue was labeled by blue box, where the number of overlapped MaPR indicated. Cancer types from validation datasets were subscript with a ’1’. (B) Overlap of MaPR of each cancer among gene sets reported with known functions. Each cell represents the percentage of gene sets with known functions that overlap with MaPRs, where cells with hypergeometric test P-value < 0.05 were boxed. Cancer genes are the union of cancer hallmark genes (Yize Li et al Cell 023) and genes from the Cancer Gene Census of Cosmic. The 2,863 drug target proteins classified into five tiers were from Savage et al Cell 2024. (C) Pagerank quantile distribution for MaPRs within or not within RBP (upper) or HKG (bottom). Significant differences based on P-value calculated by one-sided Wilcoxon test were denoted by asterisks. (D) Pruned BRCA post-transcriptional network showing protein nodes colored by MaPR and RBP overlap (upper) or by MaPR and HKG overlap (bottom). Node border thickness corresponds to PageRank centrality scores of protein nodes in the unpruned network. Abbreviations: RBP: RNA binding protein; HKG: Housekeeping Gene; TF: transcription factor.

To provide functional contexts of MaPRs, we curated multiple gene sets associated with known processes in the regulation of protein abundance (**Table S2**), including (1) modulation of protein translation, such as RNA binding proteins (RBPs)^27–29^, ribosome proteins, spliceosome proteins, and proteins involved in transportation (nuclear pore complexes), (2) modulation of protein half-life or degradation, such as autophagy regulation, proteasomes and ubiquitin proteins. As shown on **Fig. 2B and S2D**, MaPRs consistently demonstrated significant enrichment (Wilcoxon test P-value < 0.01) in processes that mediate translation rate across cancer types, but are rarely significantly enriched in processes modulating protein half-life or degradation. These results suggest a relatively strong impact of translation on overall protein abundance, especially from RBPs. As expected, we observed the lack of enrichment of MaPRs among transcription factors (TFs)^30^ (**Fig. 2B and S2D**), indicating the successful removal of transcriptional regulators by the MaPR pipeline. Housekeeping genes (HKGs) exhibited significant overlap with MaPRs among most cancer types (**Fig. 2B and S2D**), suggesting the essentiality of MaPRs in maintaining cellular fitness. MaPR were also enriched in cancer related gene and known/potential drug targets among various cancers (**Fig. 2B and S2D**), implying its potential for cancer therapy.

To illustrate these findings, we visualized the cancer-specific post-transcriptional networks derived from the MaPR pipeline to demonstrate these findings. Due to high network density, each post-transcriptional network was first pruned before visualization (**Methods**). The pruned BRCA post-transcriptional network shows MaPRs, particularly those that are RBP and HKGs, localized at the center of the network (**Fig. 2D)**. Additonally, a higher PageRank centrality score were systematically observed among RBP MaPRs or HKG MaPRs across cancers (**Fig. 2C**). These observations reinforce MaPRs as hub proteins and highlight their central regulatory roles in post-transcriptional processes.

### Post-transcriptional regulators with therapeutic potential

Analysis across 10 cancer cohorts enabled us to identify MaPRs that are cancer-specific as well as those shared across multiple cancer types. As shown in **Fig. 3A**, the majority of MaPRs (57.98%, 2375/4096) were identified in only one cancer type. In particular, GBM MaPRs accounted for the largest proportion of cancer-specific MaPRs (32.6%, 775/2375) (**Fig. 3B**), highlighting brain-specific proteome regulations that align with tissue-specific gene expression and transcript usage patterns observed within the brain^31^. In contrast, only a small proportion of MaPRs (5.42%, 222/4096) were observed in the majority of cancer types (>5 cancers). Based on the number of cancer types identified, all MaPRs were categorized into three classes: cancer-specific MaPR, moderate MaPR and pan-cancer MaPR (**Fig. 3A**).

**Figure 3.**
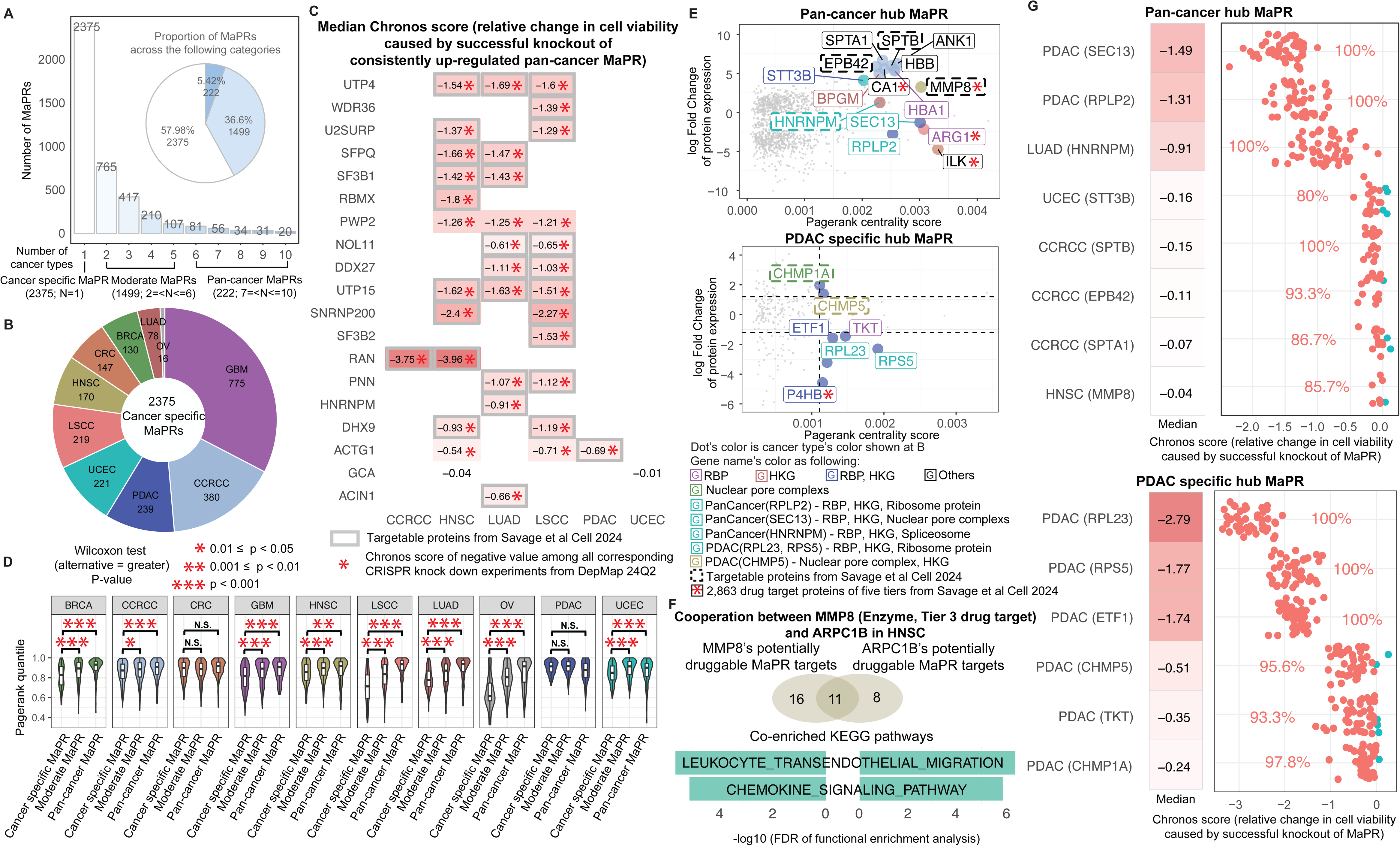
Post-transcriptional regulators and their therapeutic potential. (A) The bar plot depicts the number of proteins identified as MaPR in different numbers of cancer types. MaPRs were classified into three categories based on the number of cancer types: cancer-specific MaPR, moderate MaPR, and pan-cancer MaPR. (B) The number of cancer-specific MaPRs across cancers. (C) Median Chronos score (relative change in cell viability caused by successful CRISPR knockout) of consistently up-regulated pan-cancer MaPR in cell lines of the corresponding lineages from DepMap 24Q2. A negative value of Chronos score indicates a loss of cancer cell viability. We only displayed MaPRs with DepMap dataset available that also exhibite up-regulated protein abundance in a specific cancer type. The gene-cancer pairs supported by up-regulated expression in tumor and reduced cell viability upon CRISPR knock down in Savage et al Cell 2024 was grey boxed. The gene-cancer pairs with negative value among all corresponding CRISPR knock down experiments from DepMap 24Q2 was denoted by an asterisk. (D) Pagerank quantile distribution for MaPRs of each cancer belonging to three MaPR categories, separately. Significant differences based on P-value calculated by one-sided Wilcoxon test were denoted by asterisks. (E) Differential expression and PageRank centrality of MaPRs. In the upper panel, pan-cancer hub MaPRs, defined with top 0.2% Pagerank centrality that showed significant dysregulated protein expression in at least one cancer, are labeled. The lower panel shows PDAC-specific hub MaPRs, defined with top 2% Pagerank centrality showing significant dysregulated protein expression in PDAC. The protein name color stands for gene sets with known functions while dot color indicates the cancer type where the MaPR was identified. The X-axis represents the Pagerank centrality score and the y-axis denotes the log Fold Change of dysregulated protein expression. The MaPR supported by up-regulation in tumor and reduced cell viability upon CRISPR knock down was dash boxed. MaPRs belonging to 2,863 drug target proteins of five tiers defined by Savage et al. 2024 were denoted with asterisk. (F) Cooperation analysis between MMP8, an enzyme of Tier 3 drug target, and ARPC1B in HNSC. Tier 3 drug targets were inhibited by drugs considered investigational or experimental. The Venn diagram shows the overlap between their potentially druggable MaPR targets while the barplot shows their co-enriched KEGG pathways. Potentially druggable MaPR targets of one MaPR were defined as its targets overlapped with potentially druggable proteins that also carry positive semi-partial correlation with the MaPR (Method). (G) Median Chronos score (relative change in cell viability following the successful CRISPR knockout) of pan-cancer hub MaPR (upper) or PDAC-specific hub MaPR (lower) in cancer cell line of corresponding lineage from DepMap 24Q2. The percentage of corresponding CRISPR knock down experiments with negative Chronos score is labelled on the points plot.

Given the close association between abnormal protein expression and tumor progression, we identified MaPRs that show dysregulated protein expression by conducting a differential expression analysis (BH-corrected P-value < 0.05 & |log Fold Change| > 1.2) among the 8 cancer types with available patient-matched adjacent normal samples (**Fig. S3A and S3B, Table S1, Methods**). Almost all pan-cancer MaPRs (221/222) displayed significant dysregulated protein expression in at least one cancer, whereas cancer-specific and moderate MaPRs were less frequently differentially expressed (**Fig. S3C**). MaPRs were split into four classes considering the consistency of dysregulated protein expression direction across cancer types: not dysregulated, up-regulated, down- regulated and mixed dysregulated (**Method**). A large portion of pan-cancer MaPRs (29.73%, 66/222) displayed down-regulated expression in all cancers, while a smaller fraction (9.01%, 20/222) showed up-regulated expression in all cancers (**Fig. S3D and S3E**). All up-regulated cancer-MaPR pairs displayed strong genetic dependency as shown by their negative chronos score, which indiciates that knockout of these MaPRs decrease cancer cell viability, in the corresponding lineages from Cancer Dependency Map (DepMap)^23^, with the majority (81.1%, 30/37) corroborated by evidence from Savage et al^3^ (**Method and Fig. 3C ans S3E**). The comprehensive list of MaPRs corroborated by Savage et al^3^ with all levels of annotated evidence across seven cancer types is provided at **Table S1**. RAN, an up-regulated MaPR, exhibited the strongest cancer cell genetic dependency in CCRCC and HNSC (**Fig. 3C)**. This could be explained by its involvement in multiple cancer pathways including PIK3-related, KRAS-related and mitosis pathways^32–34^. These analyses highlight dysregulated MaPRs that are critical in cancer cell growth and progression.

Previous transcriptomics research indicates that hub genes with higher centrality scores are more influential in regulatory networks, but the importance of hub proteins in post-transcriptional network remains to be characterized. Across post-transcriptional networks of most cancer types, non-cancer-specific MaPRs often exhibited higher centrality scores (**Fig. 3D**). To identify MaPRs with highest impact, we selected the top 0.2% of pan-cancer MaPRs ranked by PageRank centrality score, that also showed differential protein expression and identified 15 hub pan-cancer MaPRs, including 6 MaPRs involved in processes known to modulate translation rate and 4 MaPRs exhibiting up-regulated expression in tumors and cancer cell genetic dependency corroborated by Savage et al^3^ (**Fig. 3E**). HNRNPM, a spliceosome component RBP responsible for RNA splicing and processing, emerged as an up-regulated hub MaPR in the LUAD patient tumor cohort that also showed strong cancer cell genetic dependency in LUAD^23^ (**Fig. 3G**). In HNSC, MMP8 was an up-regulated potentially druggable enzyme in tumors and showed cancer cell genetic dependency^3^, which was also reported up-regulated in HNSC patient serums^35^. We further investigated the therapeutic potential of MMP8 MaPR networks by combining evidences from druggability compiled by Savage et al^3^, our DEP analysis, and MaPR post-regulatory networks (**Method**). In our HNSC cohort analysis, MMP8 was an up-regulated hub MaPR with its associated potentially druggable MaPR targets enriched in two immune-related pathways: leukocyte transendothelial migration and chemokine signaling pathway. Interestingly, cooperation analysis identified another MaPR, ARPC1B, whose potentially druggable MaPR targets were also enriched in the same two pathways in HNSC (**Method**). This finding was supported by previous studies identifying ARPC1B as promising target for tumor immunotherapy and prognostic biomarker in various cancers, including HNSC^36,37^. We further identified additional MaPR pairs, including SLC12A7-NCEH whose potentially druggable MaPR targets were enriched in the same six pathways in PDAC and WDR5-AQR in four pathways in LSCC (**Fig. S4A and S4B and Method**). Notably, 14 MaPRs’ potentially druggable MaPR targets were enriched the same two pathways, N glycan biosynthesis and protein export, as another potentially druggable enzyme MaPR, FKBP11, in LSCC (**Fig. S4C and Method**).

We additionally identified dysregulated cancer-specific hub MaPRs within the top 2% of PageRank centrality scores (**Fig. 3E and S4E**). In PDAC, we identified MaPRs with cancer cell genetic dependency, including RPS5 and RPL23, two tumor down-regulated ribosome proteins that participates in translation initiation, CHMP5, a housekeeping gene that regulates late endosome function^38,39^ and CHMP1A, a nuclear pore complex protein (**Fig. 3E and 3F**). Up-regulated expression in tumors and cancer cell genetic dependency of the two MaPRs, CHMP5 and CHMP1A, were also observed in Savage et al^3^ (**Table S1**). Moreover, we identified MaPRs with therapeutic potential shared across cancer types, including DDX21 (MaPR in BRCA and LUAD), RSL1D1 (MaPR in BRCA and LUAD) and SMC2 (MaPR in LUAD, HNSC and LSCC). These three MaPRs showed cancer cell genetic dependency^23^ (**Fig. S4D**) and our recent knockdown experiments of these three MaPRs reduced cancer cell survival^40^. Overall, these analyses highlight key MaPRs that are central to post-transcriptional regulation networks and with therapeutic potential in multiple cancer types.

### Pathways enriched for MaPRs and post-transcriptional regulatory networks

To examine how MaPRs may perburb pathways underlying tumorigesis, we performed a functional enrichment analysis (**Methods**). We selected the top 30% of MaPRs ranked by Pagerank centrality score and calculated their statistical overrepresentation in KEGG pathways using the hypergeomeric test (BH-corrected P-value < 0.05, **Fig. 4A and Table S2**). Diverse pathways were perturbed by these central MaPRs, including post-transcriptional processes, such as ribosome, splicesome, and protein transports, as well as tumor-related pathways, such as oxidative phosphorylation and metabolism. Pathways for multiple neurodegenerative diseases also showed significant enrichment for MaPRs, possibly due to the involvement of peptide processing and misaggregation in these ontologies (**Fig. 4A and 4B**).

**Figure 4.**
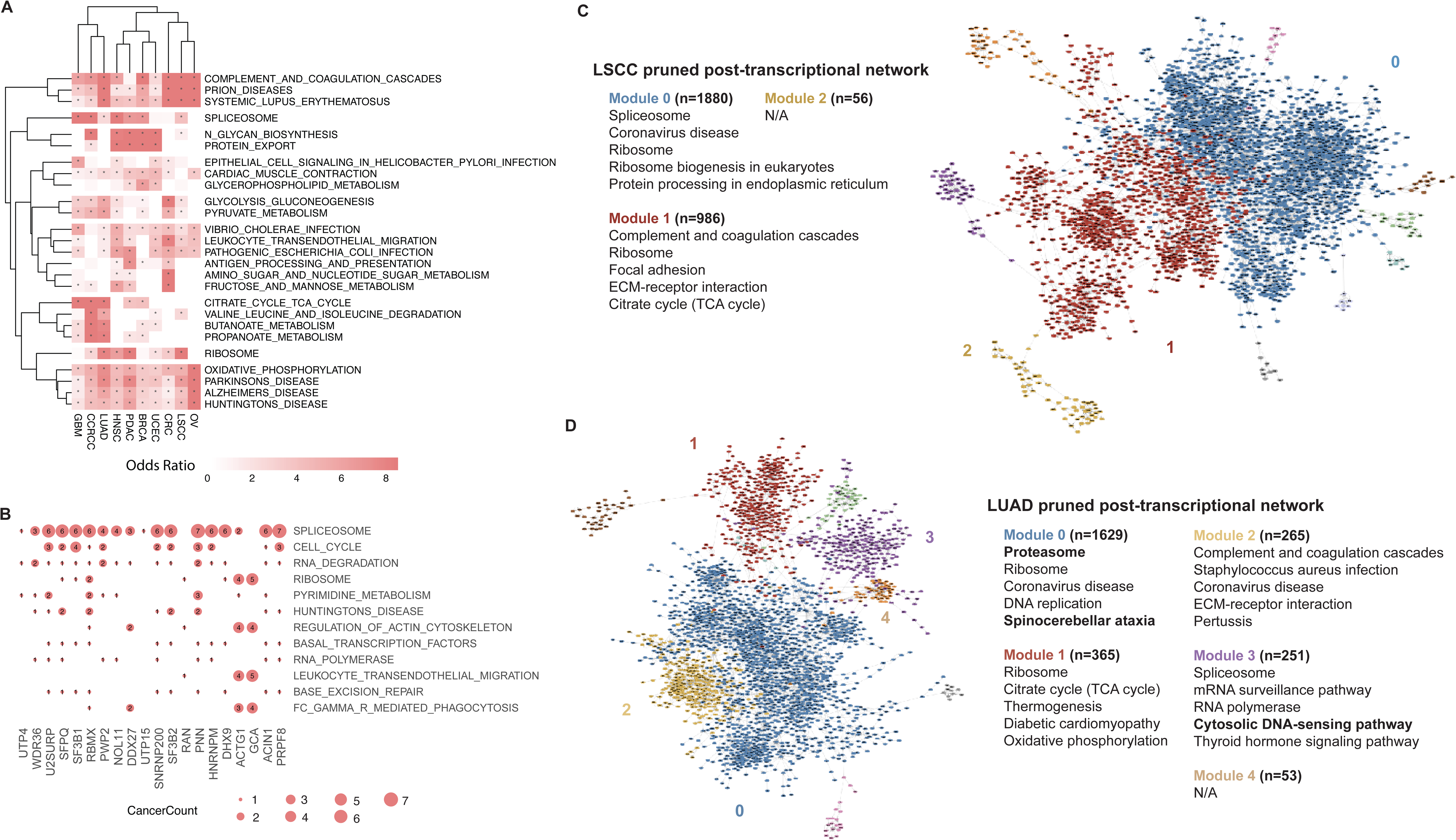
**Pathway enrichment reveals biological processes modulated by MaPR proteins and targets**. (A) Top pathways significantly enriched by MaPRs with top 30% Pagerank score in each cancer. The color of each cell indicates the odds ratio of the pathway enrichment, where the odds ratios that were greater than 1 are denoted by an asterisk. In the plotting scale, odds ratio greater than 1.5 times the interquartile range above the third quartile were reduced to this value. (B) Pathway enrichment of selected MaPR’s targets, where each cell represents the number of cancer types where an enriched pathway. These MaPRs were chosen from pan-cancer MaPR that shows consistently up-regulated expression across cancer types. (C-D) Pruned post-transcriptional networks in LSCC and LUAD, respectively. Protein nodes are colored by protein module membership found using the greedy modularity maximization algorithm. Proteins within each module, containing at least 50 nodes, were subject to enrichment analysis, and the top 5 significantly enriched KEGG pathways are shown. Significantly enriched KEGG pathways that are both in the top 5 and cancer-specific (only significantly enriched by either LUAD or LSCC, not both) are in bold font.

We also performed functional enrichment analysis of all the predicted targets of each MaPR in each cancer. Despite pervasive pathway enrichment by MaPRs’ targets, target-enriched pathways varied across cancers. The top enriched pathways overall includes spliceosome (**Fig. S5B**), which was also prominently enriched by consistently up- regulated pan-cancer MaPRs (**Fig. 4B**). Given previous studies highlighting the discrepancy of translation efficiency for diffent transcript isoforms^41,42^ orchestrated by the spliceosome, this finding underscores the influence of MaPRs on protein abundance via alternative splicing. Additionally, our analysis captured pathway patterns affected by critical MaPRs. For example, targets of the tumor up-regulated MaPR PRPF8 were enriched for the spliceosome pathway across all the seven cancers where it was identified as a MaPR (**Fig. 4B**). Two up-regulated MaPRs, ACTG1 and GCA, were similarly enriched for targets in four pathways, including ribosome (**Fig. 4B**). The top 10 MaPRs showing the highest numbers of target-enriched pathways were associated with the electron transport chain and ATP production (**Fig. S5A**) and their targets were showed the most significant enrichment for oxidative phosphorylation (**Fig. S5C)**, implying roles of MaPRs in energy production.

Comparing post-transcriptional networks and enriched pathways in LSCC and LUAD, the top MaPRs for each cancer type shared enrichment in essential pathways of ribosome, spliceosome, oxidative phosphorylation, complement and coagulation cascades and leukocyte transendothelial migration. Meanwhile, these two lung cancer types also displayed cancer-specific enrichment in several metabolism pathways (**Fig. 4A**). Similar to the top MaPRs enrichment analysis, the targets of LSCC/LUAD’s 331 shared MaPRs were enriched in essential pathways while distinct metabolism pathways were enriched by the targets of the cancer-specific MaPRs (**Fig. S5D and S5E**). Butanoate, pyruvate and propanoate metabolism were enriched in LUAD (**Fig. S5F**), while sugar, purine, and pyrimidine metabolism were enriched in LSCC (**Fig. S5G**). **Figure 4C and 4D** provides visualization of the pruned post-transcriptional networks for LSCC and LUAD. KEGG pathway enrichment analysis were conducted^43^ within each of the modules that included at least 50 proteins (**Methods**, **Table S3)**. Both cancer types shared module enrichment for terms such as ribosome, spliceosome, protein processing in the endoplasmic reticulum, oxidative phosphorylation, and complement and coagulation cascades **(Fig. 4C and 4D, Table S3)**. However, both cancer types displayed module enrichment for unique KEGG pathways as well (**Table S3)**. For example, LSCC module 0 displayed unique enrichment for KEGG terms such as NF-kappa B signaling pathway, hematopoeic cell lineage, cellular senescence, ubiquitin mediated proteolysis, and galactose metabolism. On the other hand, LUAD module 0 displayed unique enrichment for KEGG terms cuh as proteosome, spinocerebellar ataxia, Parkinson Disease, and Alzherimer’s Disease. These results demonstrate how MaPR-mediated post-transcriptional regulation may delineate the unique features and mechanisms defining tissue-specific cancer types.

### Abundance of MaPRs predict proteome subtypes across cancers

We hypothesized that if MaPRs play a central role in regulating post-transcriptional processes, then MaPR levels, compared to other proteins, can reasonably predict and define the overall proteome observed in tumor samples. To test this hypothesis, we first identified proteome subtypes by clustering samples within each cancer type based on their whole proteomic profiles (**Fig. 5A, 5B and S5H**) (**Methods**). Next, using protein levels as a proxy for activity, we trained random forest classifiers to predict proteomic subtype membership with randomly selected (n=100 or 10) MaPRs and same number of other proteins for 1,000 permutations in each cancer type^44^ (**Methods**). Each model’s performance was assessed using 5-fold cross validation and the mean AUC ROC was calculated. As the number of features decreased from 100 to 10, MaPRs generally showed increased predictability of proteomic subtype compared to same number of non- MaPRs across all cancer types (**Fig. 5C, Table S4**). For example, randomly selected 100 BRCA MaPRs and 100 other proteins both predict subtype membership with high performance (Mean AUC ROC MaPRs: 0.973, Mean AUC ROC Other proteins: 0.988, P-value: 3.56e-148, Mann-Whitney U test^45^), suggesting that 100 proteins can reasonably reconstitute overall proteome subtypes. However, when the number of features dropped to 10, randomly selected BRCA MaPRs performed significantly better than other proteins in predicting subtype membership (Mean AUC ROC MaPRs: 0.882, Mean AUC ROC non- MaPRs: 0.865, P-value: 9.42e-09, Mann-Whitney U test^45^). In CRC, the same trend was observed where 100 randomly selected MaPRs and 100 other proteins both perform well at predicting subtype membership (Mean AUC ROC MaPRs: 0.952, Mean AUC ROC Other proteins: 0.948, P-value: 3.69e-3, Mann-Whitney U test^45^). However, a decrease in performance was observed in the predictive ability of other proteins when the number of features dropped to 10 (Mean AUC ROC MaPRs: 0.916, Mean AUC ROC Other proteins: 0.825, P-value: 7.82e-208, Mann-Whitney U test^45^). This trend was observed across all 10 cancer types and suggests that individual MaPRs hold more information reguarding the cancer proteome than other proteins. Overall, these findings support our hypothesis that MaPRs are playing a central role in regulating protein abundance.

**Figure 5.**
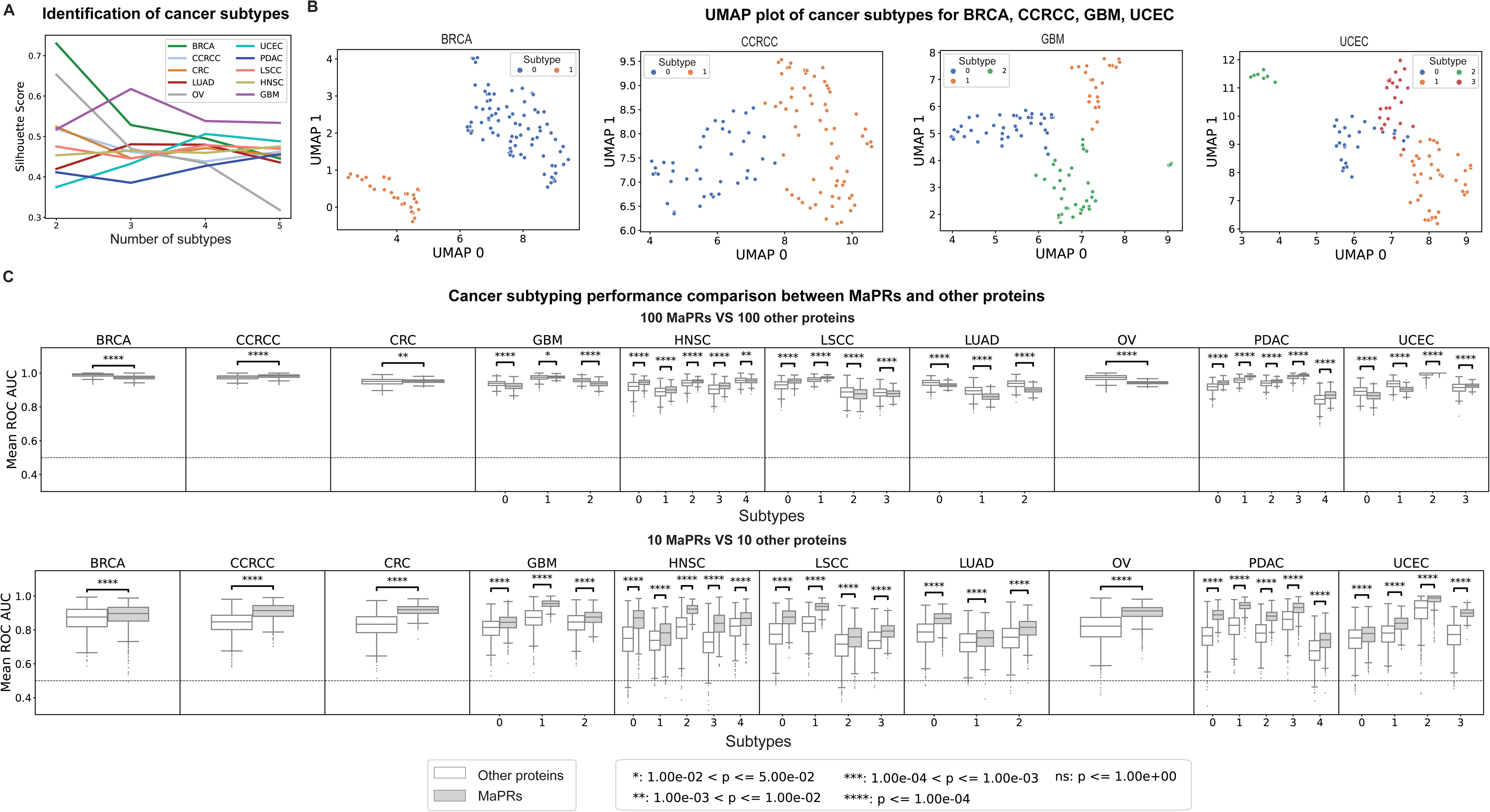
Performance of MaPRs vs other proteins in predicting cancer subtype. (A) Automatic identification of the optimal number of subtypes for each cancer type. UMAP was applied to the proteomics data and K-Means clustering was applied to the first 3 UMAP components. Silhouette scores were calculated for 2-5 cancer subtypes. (B) UMAP visualization of subtype membership for an optimal number of subtypes found for BRCA, CCRCC, GBM, and UCEC. (C) Predicting tumor proteome subtypes based on 100/10 MaPRs vs. non-MaPR proteins. The protein abundances of a randomly selected set of 100 (top row) and 10 (bottom row) MaPRs/other proteins were used as features to train a random forest classifier to predict cancer subtype membership. For each round of training (1,000 permutations), 5-fold cross-validation was performed and the mean ROC AUC was calculated (boxplots). A Mann-Whitney U test was performed to assess the statistical difference between ROC AUCs for MaPRs and other proteins.

### Validation of MaPR targets based on eCLIP and knockdown experiments

To validate predicted MaPRs, we obtained RNA binding targets of RBPs measured by the ENCODE eCLIP experiments^46^, and knockdown RNA-seq data across two cell lines, K562 and HepG2^47^ (**Methods**). Among the 139 RBPs that had data available for both eCLIP and knockdown RNA-seq within identical cell line, 85 RBPs were identified as MaPR in at least one cancer. MaPR RBPs showed significantly higher target validation rates compared to non-MaPR RBPs based on eCLIP binding of the targets’ RNA, as well as the targets’ differential mRNA expression/splicing upon knocking-down the corresponding MaPR (**Fig. 6A**). Combining eCLIP and knockdown differential expression to validate MaPR targets revealed 7 MaPRs with significant enrichment of targets being validated, among which 6 MaPRs were splicesome factors belonging to three protein families associated with pre-mRNA splicing: heterogeneous nuclear ribonucleoproteins (hnRNPs), RNA-binding motif (RBM) proteins and serine/arginine (SR)-rich proteins (**Fig. 6B**). This is expected as other MaPRs modulating only protein exports/processing were less likely to affect differential mRNA expression, whereas splicesome MaPRs modulate both transcription and translation processes that could be tightly linked. These results align with the splicesome’s central role in differential transcript usage/generation and modulation of translation rate, thereby influencing protein abundance^41,42^.

**Figure 6.**
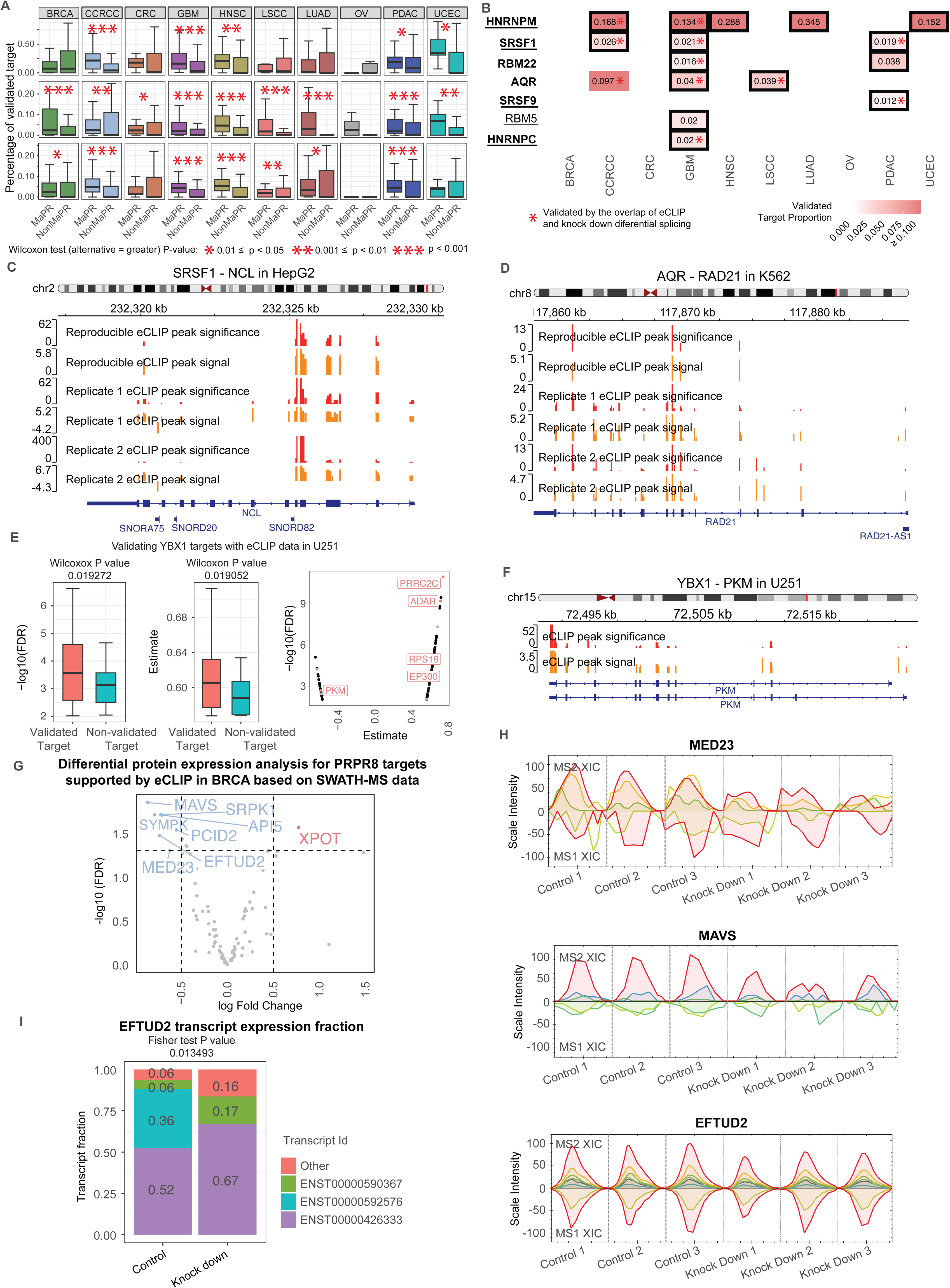
Experimental validation of MaPR-target regulation based on eCLIP and knockdown datasets. (A) Percentage of predicted targets of MaPR and other Non- MaPRs validated by eCLIP (top), knockdown differential expression (middle, |log2Fold Change| > 1.2 & FDR < 0.05) and knockdown differential splicing (bottom). Significant differences based on P-value calculated by one-sided Wilcoxon test were denoted by asterisks. (B) MaPR targets validated by both eCLIP binding of MaPRs and differential gene expression upon MaPR knockdowns. Each cell color stands for the percentage of predicted targets validated by the overlap of eCLIP and knock-down differential expression (|log2Fold Change| > 1.2 & FDR < 0.05) in the same cell line. Only significantly validated cells are shown in red and respective validation rates. Only protein predicted as MaPR in a specific cancer is boxed. MaPRs that belong to the spliceosome pathway are bolded, and MaPRs that are housekeeping genes are underlined. Cells significantly supported by the overlap of eCLIP and knockdown differential splicing in the same cell line were denoted by an asterisk. (C) eCLIP RNA binding peaks of SRSF1 on NCL in HepG2 cell line. (D) eCLIP RNA binding peaks of AQR on RAD21 in the K562 cell line. (E) Comparison of –log10(FDR) or estimated coefficients among post-transcriptional regulatory network between eCLIP validated targets and non-validated targets of YBX1 (left and middle) in GBM. Estimated coefficients vs –log10(FDR) from the post- transcriptional regulatory network for eCLIP validated targets (right) in GBM. (F) eCLIP RNA binding peaks of YBX1 on PKM in GBM cell line of U251. For all the eCLIP peak region visualization figures, the red bar represents the significance of eCLIP peaks (– log10 of P-value) and the orange bar represents a signal of eCLIP peaks (log2 of fold change). All significant differential splicing defined as |IncLevelDifference| > 0.05 & FDR < 0.1 & P-value < 0.05. (G) Differential protein expression analysis upon PRPF8 knock down in Cal51 cell line for the 105 targets supported by eCLIP in BRCA. (H) SWATH-MS protein expresion upon PRPF8 knock down in Cal51 cell line shown by XIC graph for three PRPF8 targets: MED23, MAVS, and EFTUD2. XIC graph used the extracted ion chromatography to measure protein abundance quantifications in three biologically independent replicates (control-siRNA-treated and PRPF8-depleted). For each XIC graph, the upper panel of peak graphs represents the MS2 level ion traces while the bottom panel of peak graphs represents the MS1 level ion traces. (I) Expression fraction for the EFTUD2 transcripts whose fraction difference is> 0.1.

We further examined the RNA binding of these MaPRs to their targets’ RNA. eCLIP data showed the extensive binding of the GBM MaPR SRSF1—alternative splicing factor 1 known to be associated with with various cancers^48^—to a substantial portion of the gene body of one of its predicted targets, NCL (**Fig. 6C, Table S5**). NCL displayed significant differential expression upon SRSF1 knockdown in HepG2 (log2FC = 1.51, FDR = 1.19e- 05). Additional MaPR-predicted targets of SRSF1—ZEB2 and SRSF5 showed differential splicing upon SRSF1 knockdown in K562 **(Fig. S6B, S6C and Table S5**). As another example, the binding of GBM MaPR AQR to its target RAD21 was validated by eCLIP data, and RAD21 showed significant differential expression and splicing upon AQR knockdown in K562 (**Fig. 6D and Table S5**). Another AQR target U2AF2 also showed significant changes in both expression and splicing upon AQR knockdown in K562 (**Fig. S6D and Table S5**).

In addition to ENCODE eCLIP data, we obtained an independent eCLIP dataset of the U251 glioblastoma cell line^49^ to validate RNA binding of YBX1, a protein we identified as a MaPR in GBM. The predicted targets of YBX1 in GBM were significantly validated by the eCLIP data (Hypergeometric test P-value < 0.05) (**Table S5**). We observed that the gene bodies of the majority of YBX1 predicted targets (81.7%, 125 /153) overlapped with at least one significant eCLIP peak of YBX1 binding (P-value < 0.05). These 125 validated targets carried higher coefficients and lower FDR value in our MaPR-derived post-transcriptional regulatory network in GBM (**Fig. 6E**). Notably, the most significant eCLIP peak target identified was PKM, a gene associated with metabolism and prognosis in cancer^50,51^ (**Fig. 6F**). Additionally, several other targets associated with multiple cancers were validated, including RPS19, a spliceosome pathway factor, and EP300 (**Fig. S6E and S6F**).

Lastly, we reprocessed and anazyed a experimental proteogenomic dataset^52^ we previously generated upon delepting a key MaPR identified herein, PRPF8, in Cal51 breast epithelial-like cells to assess the resulting transcriptomic and proteomic changes quantified by RNA-seq and SWATH-MS (**Methods**). Based on the BRCA primary tumor cohort used herein, our MaPR pipeline identified that majority of predicted targets of PRPF8 (80.2%, 105/131) were supported by the eCLIP data from ENCODE (**Fig. S6A**). These targets significantly overlaped with differentially-expressed proteins upon PRPF8 knockdown in this experimental data (8 overlapped proteins, Hypergeometric test P-value = 2.08E-4, **Fig. 6G**), including MED23, reported to activate tumor cell invasion and metastatis^53^, and MAVS, whcich is associated with immune regulation^54^ (**Fig. 6H**). Seven of the differentially expressed target proteins were down-regulated upon PRPF8 knockdown (**Fig. 6H and S6G**), which is aligned with their positive semi-partial correlation with PRPF8 in the MaPR-derived post-transcriptional regulatory network of the BRCA primary tumor cohort. Interestingly, EFTUD2, a spliceosome component, showed significant differential expression at the protein level but not at mRNA level upon PRPF8 knockdown (**Fig. 6H**). This could be explained by its varied use of different mRNA transcripts that may have affected translation and eventually, protein abundance (**Fig. 6I**). Collectively, these results demonstrate that our MaPRs inferred from human tumor cohorts are validated by experimental data of RBP MaPR binding’s to target RNAs and expression changes of their predicted targets upon MaPR knockdown.

## Discussion

Protein abundance in biological systems is determined not only by mRNA expression, but also by multiple post-transcriptional processes. Our study elucidates the critical roles of Master Protein abundance Regulators (MaPRs) in post-transcriptional regulation networks across various cancer types, providing a new resource into the complex landscape of protein abundance regulation. By integrating high-throughput proteomic and transcriptomic data from ten cancer cohorts, we developed a robust computational pipeline to identify MaPRs. The identification of 232 to 1,394 MaPRs per cancer type, mediating a substantial proportion of the post-transcriptional regulatory landscape, underscores the extensive influence of these regulators that were understudied by previous system biology approaches focusing on DNA/RNA-level analyses.

Pathway enrichment analyses of MaPRs and their targets reveals the extensive impact of MaPRs on key post-transcriptional processes and tumor-related pathways. The enrichment of pathways associated with ribosome, RBPs, spliceosome, and oxidative phosphorylation across multiple cancers aligns with the known roles of these pathways in protein synthesis and energy production^1^, which are essential for cancer cell proliferation and survival. Surprisingly, MaPRs were not enriched for gene sets associated with modulation of protein’s half life and degradation, suggesting they are relatively weaker regulators of the overall proteome abundances compared to RBPs. The cancer-specific enrichment of metabolism pathways further highlights the distinct metabolic reprogramming that occurs in different cancer types^55^, a hallmark of cancer that presents opportunities for targeted therapies.

Identified MaPRs are largely cancer-specific, with the majority found in only one cancer type. This observation underscores the tissue-specific nature of post-transcriptional regulation, particularly in the context of proteome regulations specific to brain tissues, as evidenced by the high proportion of MaPRs in glioblastoma multiforme (GBM). In contrast, a small subset of MaPRs exhibit pan-cancer dysregulation, pointing to their potential role as universal regulators in tumorigenesis. The majority of pan-cancer MaPRs showed significant dysregulation, predominantly exhibiting down-regulated expression across cancers, which is consistent with their involvement in pathways crucial for maintaining cellular homeostasis and proliferation. Notably, the identification of genetic dependency in up-regulated MaPRs across multiple cancer types, as corroborated by data from the Cancer Dependency Map^23^, highlights their essential role in cancer cell survival and their potential as therapeutic targets. For example, we identified hub MaPRs that were up- regulated in tumors and show strong cancer cell genetic dependency, such as HNRNPM in LUAD and MMP8 in HNSC, which already has investigational or experimental drugs^3^.

Our hypothesis that MaPRs play a central role in defining proteome subtypes is supported by the strong predictive power of MaPRs in classifying proteomic subtypes across cancer types. The ability of using just ten MaPRs to predict proteomic subtypes with high accuracy, particularly when compared to non-MaPR proteins, suggests that MaPRs encapsulate key information about the cancer proteome and could serve as biomarkers for molecular subtyping and cancer prognosis.

The MaPR pipeline constructs post-transcriptional regulartory networks using quantitative proteogenomic profiles to identify tissue-specific regulatory patterns. This approach complements sequence-based deep learning (DL) models, which have shown excellent performance in predicting RBP and non-coding RNA (ncRNA) target binding, but have been trained on in vitro experimental data and ignore tissue contexts. For example, DeepCLIP^56^ and RBPNet^57^ focus on using sequence-based approaches to predict RBP targets. These predictions are complementary and may be combined to achieve better understanding of post-transcriptional regulations. Furthermore, ongoing developments of technologies surveying “translatomics”^58^, including polysome profiling, ribo-seq, trap-seq, proximity-specific ribosome profiling, rnc-seq, tcp-seq, qti-seq and scRibo-seq, will also provide data to be cross-validated the MaPR post-transcriptional regulartory networks. The validation of MaPR targets using eCLIP and knockdown experiments provides robust experimental evidence supporting the functional relevance of MaPRs in post-transcriptional regulation. The independent validation of MaPR binding in glioblastoma cell line, as well as pre/post-knockdown differential protein expression of MaPR targets in breast cancer cells, further strengthen the credibility of the human cohort-derived MaPR networks and validates the functional and proteome consequence of targeting MaPRs.

Our study has several limitations. First, the identification of MaPRs is based on computational predictions and statistical correlations, which, although supported by available experimental data, require further experimental validation to confirm their functional roles. Second, the variability in data quality, sample size, and proteomic data coverage may introduce biases across different cancer cohorts, and we envision the completeness and robustness of identified MaPRs will improve over time given the rapid rise in high-quality proteogenomic cohorts. Lastly, while our study identifies key MaPRs, it does not investigate their interactions with mutations or other biomolecules such as non-coding RNAs that may also participate in post-transcriptional regulation. Addressing these limitations in future research will be crucial for fully elucidating the role of MaPRs across cancer types.

In conclusion, the identification of MaPRs reveals key regulators of post-transcriptional processes in cancer. The validation of MaPR-target relationships using experimental RNA binding and knockdown data provides robust evidence supporting the regulatory networks generated by our new computational tool. Our findings not only elucidate the complex regulatory networks governing protein abundance, but also highlight the potential of MaPRs as biomarkers and therapeutic targets. This study provides a resource of the catalog of MaPRs across ten cancer types and a robust MaPR computational tool for proteogenomic cohorts, laying the groundwork for future research aimed at unraveling the intricate post-transcriptional regulatory mechanisms that shape cancer and cell biology.

## METHODS

### Paired protein/mRNA expression profiles across 10 cancer cohorts

Genome-wide paired protein and mRNA expression profiles of 1,047 samples across 10 cancer cohorts were downloaded from CPTAC (https://portal.gdc.cancer.gov/,). The protein expression is a relative quantification achieved by MS technology. Normalization of Median Absolute Deviation (MAD) is performed for protein expression of each sample within a specific cancer cohort yielding a unit MAD. Proteins without detected expression in more than 30% samples in each cancer cohort were removed. The mRNA expression was processed through STAR-Counts pipeline and measured by Fragments per kilobase of transcript per million mapped reads upper quartile (FPKM-UQ). mRNAs showing no expression in more than 30% of samples were removed. The curated samples conferred clinical diversities of different levels, such as age and gender, except for that all samples of BRCA, OV and UCEC were female. We also collected clinical information available for each cancer, including age, gender and other aspects used in downstream differential expression analysis.

In addition, genome-wide paired protein and mRNA expression profiles of 258 independent samples across 4 cancer cohorts were downloaded from The Cancer Genome Atlas (TCGA) for validation analysis. Proteome data of OV is derived from two institutions: Pacific Northwest National Laboratory (PNNL) and Johns Hopkins University (JHU), where 21 samples were shared by both institutions. The overview of paired protein/mRNA expression profiles were summarized at **Fig. 1A** and **Table S1**.

### Construction of post-transcriptional regulatory network

We developed a computational pipeline, MaPR, to construct a post-transcriptional regulartory network independent of transcriptional regulation by simultaneously analyzing transcriptomic and proteomic data of the same cohort. The pipeline consists of the following steps: (1) Construct two undirected graphs: one graph for protein co-expression network Gp = (Vp,Ep), where Vp is the set of quantified proteins in the input dataset and Ep are significant pair-wise spearman correlations of protein expression (Benjamini-Hochberg BH corrected P-value < 0.01); and the other graph for mRNA co-expression network Gm = (Vm,Em) obtained through the same method applied to the transcriptomic data. (2) Remove co-expressed protein pairs that also show co-expression at the mRNA level, thus generating a new graph Gp-only = (Vp,Ep – Em). (3) Generate a directed graph of post-transcriptional regulation. Given protein A is a candidate protein abundance regulator and B is a protein which it regulates, we reasoned that protein A’s protein expression will be correlated with protein B’s protein expression independent of protein B’s mRNA expression. Based on the Gp-only graph, we iterated through each edge and calculated the semi-partial spearman correlation between protein A and protein B^21^:

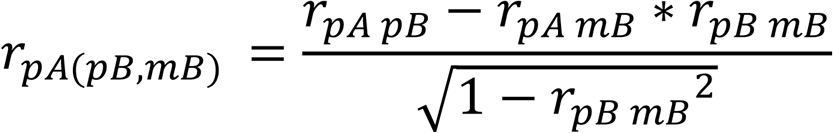

where *pA* and *pB* are the expression of protein A and protein B, respectively, *mB* is the mRNA expression of protein B, and r is spearman coefficient between variables. The final post-transcriptional regulatory network is subsequently defined with a directed graph GMaPR = (VP,EMaPR) where the directionality denotes the regulators to the regulated proteins based on significant semi-partial spearman correlations (Bonferroni corrected P-value < 0.01) corrected for the regulated protein’s mRNA level. To include more edges into consideration and retain the most significant edges as post-transcriptional regulatory network used for further MaPR prediction, the BH method and the stricter Bonferroni P-value correction method were used at the (1) and (3) step, respectively.

### Identification of MaPR

We hypothesized that, like transcription factors (TFs) that control RNA expression transcribed from DNA, there exists regulators (i.e., ribosomal-binding proteins) that mediate protein abundance translated from RNA. We used an empirical permutation-based method to identify such protein abundance regulators from GMaPR that show statistical enrichment of regulated proteins. In this procedure, all edges EMaPR were shuffled 10,000 time, and each time, the summed number of EMaPR originated from a given candidate protein abundance regulator was calculated. The significance P-value for each regulator is simply calculated as the fraction of random simulations where the originating regulator edges is larger than that observed for a given protein in GMaPR. This procedure was repeated 100 times to identify significant regulator as MaPRs and the overlapped MaPRs of repetition were selected as final predicted MaPRs.

### Differential expressed protein (DEP)

To identify differentially expressed protein, limma R package was used to perform a patient-paired (tumor and normal sample from the same patient) differential expression analysis in each cancer cohort, where we corrected any available confounding factors in that cancer cohort including demographics (age, ethnicity, race and geneder) and batch effects (sequencing center, generating date, operator and tandem mass tag TMT batch). Significant was defined as both BH-corrected P-value < 0.05 and absolute value of log Fold Change > 1.2. Specially, no enough patient matched tumor/normal samples exist in GBM and OV leading no DEP identificationin among these two cancer types (**Table S1**). We classified MaPRs into four categories considering the consistency of dys-regulated protein expression direction across cancers. Given one MaPR dysregulated in A cancers, including up-regulated in B cancers and down-regulated in C cancer, it is classified as (i) not dysregulated if A = 0; (ii) up-regulated if A > 0 & A = B & C = 0; (iii) down-regulated if A > 0 & A = C & B = 0; (iv) Mixed dys-regulated otherwise.

### Pruning, visualization, and module identification of post-transcriptional networks

In order to visualize the dense post-transcriptional networks for each cancer type, the networks were first pruned. The pruning process involves ranking all edges for each regulator using the -log10(FDR) value and preserving only the top 2 edges. The pruned networks were then visualized using Cytoscape v3.10.1^59^ using the Prefuse Force Directed Layout with default settings. Protein modules were identified using Clauset-Newman-Moore greedy modularity maximization^60^ (resolution = 0.1) using the NetworkX v3.2.1^61^ Python package. Finally, proteins without connections to the main network due to the pruning process were removed to simplify visualization.

### Centrality score of MaPR in each cancer type

Pagerank^62^ is an algorithm originally developed to rank the importance of websites during a Google search and successfully applied to address biological problems. We calculated Pagerank scores to measure the centrality of each node in the post-transcriptional network. For each cancer type, the post-transcriptional regulatory network was initiated as a directed graph using the NetworkX^61^ Python package. Proteins in the network are represented as nodes and edges connecting nodes are directional, originating from a MaPR and pointing towards its target(s). The -log(FDR) was used as edge weights due to its direct relationship with the absolute value of the Spearman correlation coefficient. The direction of all edges in graph were reversed before calculating the PageRank centrality (edges originating from targets and pointing towards their regulators).

### Functional Enrichment Analysis

We used proteins within expression profile filtering gene set associated with KEGG pathways obtained from Msigdb database^63^, resulting in 186 KEGG pathways ranging from 8 genes to 279 genes (**Table S2**). Hypergeometric test was performed to check whether gene set of interest were significantly overpresented in each KEGG pathway, where genes with expression available used as background. Enrichment significance is defined as both more than three overlapped and BH-corrected P-value < 0.05. We analyzed two gene sets of interest, including: (i) targets of each MaPR of each cancer type, (ii) top MaPRs in terms of their Pagerank score within each cancer type.

### Cancer Dependency Map (DepMaP)

To assess MaPR’s effect on cell viability, we acquired gene dependency scores from broad CRISPR experiments in cancer cell lines of corresponding lineage, sourced from current version 24Q2 of Cancer Dependency Map (DepMap)^23^. We used ‘OncotreeCode’ of DepMap model file to associate cancer cell line with our cancer types, where only two cancers need to notice that ‘GB’ classified as ‘GBM’ and ‘PAAD’ classified as ‘PDAC’ (**Table S1**). The gene dependency scores were calculated using the Chronos algorithm^64^, measuring the relative change in growth rate resulting from successful knockout of the gene. A score of 0 indicates no change in cancer cell viability, negative value indicates a loss of viability and positive value indicates a gain of viability.

### Targetable dependency driven by up-regulated expression in tumors and reduction in cell growth viability

Savage et al expanded the landscape of therapeutic targets by combinatorial analysis of CPTAC proteome dataset and CRISPR experiment from an older version 22Q2 of DepMap^3^. We obtained the list of targetable proteins with up-regulated expression in tumors of CPTAC that also reduce cell growth in DepMap from their supplementary material (Table S3). These proteins, exhibiting differential protein abundance in our analysis at same cancer cohort of CPTAC, were identified as potentially druggable proteins for further analysis across seven cancers with all dataset available (CCRCC, CRC, HNSC, LSCC, LUAD, PDAC and UCEC, **Table S1**).

### Identification of cooperative MaPR pairs with therapeutic potential in each cancer type

Cooperation between MaPR with therapeutic potential, MaPRs that belongs to potentially druggable proteins and 2,863 drug target proteins of five tiers from Savage et al^3^, and other MaPR are defined in a cancer type if they shared potentially druggable MaPR targets that are significantly enriched in at least two KEGG pathways. Potentially druggable MaPR targets of one MaPR were defined as its targets overlapped with potentially druggable proteins that also carry positive semi-partial correlation with the MaPR. Herein, the cooperation analysis were only performed on tumor up-regulated MaPR in our differential protein abundance analysis across seven cancer types.

### Machine learning to predict proteome subtype membership from MaPR protein abundance

CPTAC proteomic profiles were first preprocessed with the following steps: 1) profile selection, 2) removing features (i.e., proteins) with many missing values, 3) imputing missing values, and 4) scaling. First, CPTAC profiles with matching proteomics and transcriptomics data were retained and proteins with missing values for >= 60% of profiles were removed. Next, missing values were imputed using k-nearest neighbors (KNN) with scikit-learn’s^44^ KNNImputer function. Predicted values were determined using the 5 nearest neighbors (n_neighbors=5) while also factoring in their distances (weights=distance). Finally, features (i.e., proteins) were Z-Score normalized using scikit-learn’s^44^ StandardScaler function. After preprocessing, Uniform Manifold Approximation and Projection (UMAP) was applied to reduce the dimensionality of the protein features and the first 3 UMAP components were retained. Subsequently, K-means clustering was subsequently applied to the first 3 UMAP components for a range of 2-5 subtypes. Finally, silhouette scores were calculated and used to determine the optimal number of subtypes for each cancer type.

The aforementioned preprocessed CPTAC proteomics profiles (Steps 1-2) and proteomic subtypes for each cancer type were then used to train a machine learning classifier. For each cancer type, protein levels of n randomly selected MaPRs were used to train a random forest classifier with 5-fold cross validation to predict each sample’s subtype membership. Scikit-learn’s^44^ RandomForestClassifier was used with default hyperparameters. Samples were split into stratified folds using scikit-learn’s^44^ StratifiedKFold function to ensure balanced cluster labels across folds. Training folds were KNN imputed and Z-Score normalized independently from the testing fold before each training iteration to prevent information leakage. Model performance was assessed by calculating the mean area under the receiver operating characteristic curve (AUC ROC) across all 5 folds. For cancers with > 2 subtypes, AUCs were calculated using the one-vs-all method. Models were trained with 100 and 10 randomly selected cancer-specific MaPRs over 1000 permutations. Using the same method, other proteins (not MaPRs) were also randomly selected as features for comparison. The same random state was used for all models as well as sample stratification. To assess statistical difference between mean AUC ROCs over 1,000 permutations for MaPRs versus non-MaPRs, a Mann-Whitney U test was performed using python package statannotations^45^.

### Validation based on experimental data eCLIP data of target RNA binding

We downloaded genome-wide eCLIP reproducible RNA binding peak region of 150 human RBPs in K562 or HepG2 cell lines from ENCODE (file set accession identifier ENCSR456FVU)^47^. Then, we inferred eCLIP-based translational regulation by using bedtools *intersect* function to overlap the reproducible RNA binding peak region of each RBP with protein-coding gene body annotation from GENCODE release 19, where one or more base(s) overlapped in the same strand defined as a regulation from the RBP to the mRNA. To validate MaPR-predicted targets by eCLIP-based targets, hypergeometric test is performed for each MaPR in a specific cancer, where significance required BH corrected P-value < 0.05 and at least 3 overlapped genes. A similar procedure was performed to the eCLIP data of RBP *YBX1* in U251 cell line^49^.

### RNA-seq upon RBPs knockdown

We downloaded the batch corrected differential expression and differential splicing upon the knockdown RNA-seq of 263 human RBPs in K562 or HepG2 cell lines from ENCODE (file set accession identifier ENCSR870OLK)^47^. Significance were defined, separately, to obtain reliable target as following: |log2(fold-change)| > 1.2 and FDR < 0.05 for differential expression while |IncLevelDifference| > 0.05 & FDR < 0.1 & P-value < 0.05 for differential splicing. There were 139 RBPs with both eCLIP and knockdown RNA-seq within identical cell lines (**Table S5**). To validate MaPR-predicted targets by knockdown-based targets, hypergeometric test is performed for each MaPR in a specific cancer, where significance required BH corrected P-value < 0.05 and at least 3 overlapped genes.

### RNA-seq, SWATH-MS proteomics (sequential window acquisition of all theoretical spectra-mass spectrometry) upon PRPF8 knockdown

We obtained transcriptome and proteome upon PRPF8 knockdown in Cal51 cell lines from our previous work^52^. Protein expression was measured by SWATH-MS in three independent experiments (control-siRNA-treated and PRPF8-depleted). The proteomics dataset was reprocessed using the latest version of Data-independent acqustion (DIA) data analysis software by using Spectronaut v18.0 and default settings^65^. Differential protein analysis were performed using standard procedures in limma R package. RNA expression, measured by raw read count, was obtained from sequencing on Illumina HiSeq2000 platform in 3 control-siRNA-treated and 4 PRPF8-depleted experiments. The limma-voom method was used to normalize raw read count to better fit standard limma R package procedures for differential RNA analysis.

## Abbreviations

*BH*: Benjamini-Hochberg
CPTAC: Clinical Proteomic Tumor Analysis Consortium
CRL: Cullin Ring ubiquitin Ligase complex
DepMap: Dependency Map
DEP: Differential Expressed Protein FDR: False Discovery Rate
FPKM-UQ: Fragments Per Kilobase of transcript per Million mapped reads Upper Quartile
HKG: Housekeeping Gene
KEGG: Kyoto Encyclopedia of Genes and Genomes
MAD: Median Absolute Deviation
MaPR: Master Protein abundance Regulator
MS: Mass Spectrometry
RBP: RNA Binding Protein
TF: Transcription Factor
SWATH-MS: Sequential window acquisition of all theoretical spectra-mass spectrometry

## Data and code availability

Data used in this publication were generated by the National Cancer Institute Clinical Proteomic Tumor Analysis Consortium (CPTAC). Data for CPTAC cohorts can be found on CPTAC data portal: https://pdc.cancer.gov/. The R package for predicting MaPR was deployed at GitHub (https://github.com/Huang-lab/MaPR), which could be installed by devtools::intall_github function. eCLIP peaks were visualized by Integrative Genomics Viewer (https://igv.org/app) with default expand mode of Refseq gene tracks.

## Supporting information

Supplementary Information (include Fig S1-2 and other sections)

## Acknowledgements

The authors wish to acknowledge the participating patients and families who generously contributed to the CPTAC data. The authors thank all members of the Huang lab for constructive discussion and Dr. Yiwen Chen for providing valuable insights on analyzing GBM eCLIP data. Large language models (LLM), including those provided through GPT models, GitHub Copilot, and Amazon CodeWhisperer may have been used in the initial drafts of coding, literature review, and writing of this work. All final codes and texts have been extensively edited and verified by the authors. This work was supported in part through the computational and data resources and staff expertise provided by Scientific Computing and Data at the Icahn School of Medicine at Mount Sinai and supported by the Clinical and Translational Science Awards (CTSA) grant UL1TR004419 from the National Center for Advancing Translational Sciences. Research reported in this publication was also supported by the Office of Research Infrastructure of the National Institutes of Health under award number S10OD026880 and S10OD030463. The content is solely the responsibility of the authors and does not necessarily represent the official views of the National Institutes of Health. This work was supported by NIH NIGMS R35GM138113 and ACS RSG-22-115-01-DMC to K.H.

## Contributions

K.H. and Z.W. conceived the research, and K.H., Z.W. and M.W. designed the analysis. Z.W. performed most of the analysis. M.W. calculated the centrality score, visualized networks, and performed machine learning analysis for cancer subtyping. J.M. helped functional enrichment analysis. W.S. identified the hub proteins from protein co-expressed network. Y.L processed SWATH-MS data upon PRPF8 knock down and generated XIC graph plot. Z.W. and A.E. downloaded and processed data from CPTAC. K.H. supervised the study. All the authors read, edited, and approved the manuscript.

## Declaration of generative AI and AI-assisted technologies in the writing process

During the preparation of this work the authors used ChatGPT, Perplexity, and Claude in order to refine language and assist with editing the authors originally written content for improved readability. After using this tool/service, the authors reviewed and edited the content as needed and take full responsibility for the content of the publication.

