## Supplementary Information (include Fig S1-2 and other sections) for "Master regulators governing protein abundance across ten human cancer types"

**Supplementary Tables**

**Supplementary Table S1.** Summary of network and factors during MaPR prediction process. Tab ‘Pan-Cancer Cohort’ summarized the sample characteristics for the 10 discovery datasets of 10 cancers and 4 validation datasets of 3 cancers used in the analysis. Tab ‘MaPR’ listed predicted MaPR across the 10 discovery datasets and 4 validation datasets. Tab ‘PPI validation’ provided the count/proportion of our post-transcriptional regulatory network validated by protein-protein physical interaction from string database. Tab ‘PotentiallyDruggableProtein’ listed potentially druggable proteins for cancers with dataset available. Tab ‘DrugTargetOfFiveTiers’ listed the 1,875 proteins within our expression profile that belongs to the 2,863 drug target proteins of five tiers from Savage et al Cell 2024. Tab ‘MaPRTherapeuticPotential’ provided the overlap of MaPR with (i) potentially druggable proteins (ii) 2,863 drug target proteins of five tiers from Savage et al Cell 2024 across cancers. Tab ‘DepMapCellLineToCancer’ listed the corresponding relationship between our cancer type and ‘Oncotreecode’ of DepMap model file.

**Supplementary Table S2.** Tab ‘FunctionalGeneSet’ listed detailed information of all functional gene set used in the project. Tab ‘KEGG_genes’ showed the number of genes of each KEGG pathway. Tab ‘KEGGEnrichedByTopMaPR’ provide the detailed information of enrichment analysis for the top KEGG enriched by the top MaPRs in terms of their Pagerank centrality score in each cancer (supplementary to **Fig. 4A**).

**Supplementary Table S3.** Tab ‘LC Module Enrichment Results’ contains Enrichr links to enrichment analysis results for protein modules with at least 50 proteins identified in LUAD and LSCC. Tab ‘LUAD + LSCC’ contains KEGG terms significantly enriched (Adjusted P-value < 0.05) by modules in both LUAD and LSCC. Tab ‘LSCC Only’ contains KEGG terms significantly enriched by LSCC modules only. Tab ‘LUAD Only’ contains KEGG terms significantly enriched by LUAD modules only.

**Supplementary Table S4.** Summary of machine learning permuation results comparing AUC ROCs between randomly selected MaPRs and other proteins. Tab ’10 features’ contains summary results for 10 randomly selected MaPRs or other proteins over 1000 permutations. Tab ’100 features’ contains summary results for 100 randomly selected MaPRs or other proteins over 1000 permutations. P-values were calculated using a two-sided Mann-Whitney U test.

**Supplementary Table S5.** Tab ‘ENCODE’ detailedly described the information of eCLIP, knockdown RNA-seq from ENCODE, where knockdown differential expression used cut off of |log2Fold Change| > 1.2 and FDR < 0.05. Tab ‘eCLIP_YBX1’ provided the validation of predicted target of YBX1 by our eCLIP peaks at different significance level. Tab ‘GenesShown’ provided the annotation details about the genes at **Fig. 6 and S6**.

**Supplementary Figures**

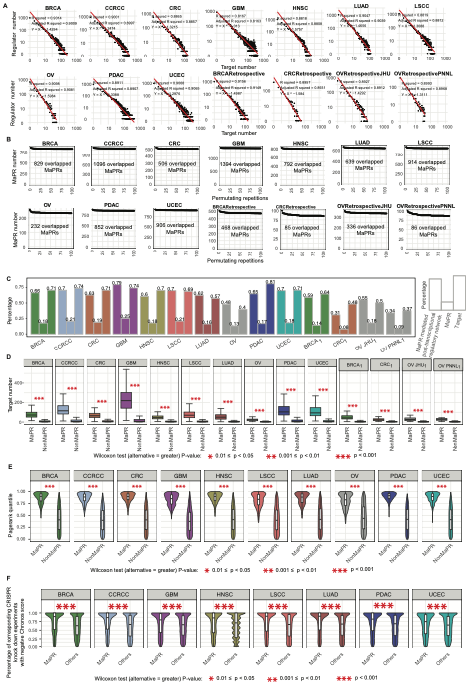

**Figure S1.** (A) The power law distribution for target number of each regulator among the post-transcriptional regulatory network across 14 datasets. X-axis denotes target number level while Y-axis represents regulator number level. (B) Overlapped MaPR count among different permutating repetitions across 14 datasets. X-axis represents the times of permutation conducted. Y-axis represents the count of overlapped MaPR of prior permutations. The overlapped MaPR count among all the 100 permutating repetitions labelled out. (C) The three columns of each cancer represent percentage of post-transcriptional regulatory network mediated by predicted MaPR, predicted MaPR and the associated target. (D) Comparison of target number between MaPR and other non-MaPR regulators. P-value calculated by Wilcoxon test (alternative = greater) for every paired target number and denoted by asterisks. Cancer type from validation dataset at (C-D) were subscript with a ’1’. (E) Pagerank quantile distribution for MaPRs and other non-MaPR regulators. P-value calculated by Wilcoxon test (alternative = greater) for every paired Pagerank score and denoted by asterisks. (F) Percentage of CRISPR knock out experiments with negative Chronos score for MaPR and other proteins in corresponding cancer line from DepMap 24Q2. Only cancer with corresponding cancer line available from DepMap 24Q2 is shown here. P-value was calculated by Wilcoxon test (alternative = greater) for every paired percentage and denoted by asterisksn in each cancer.

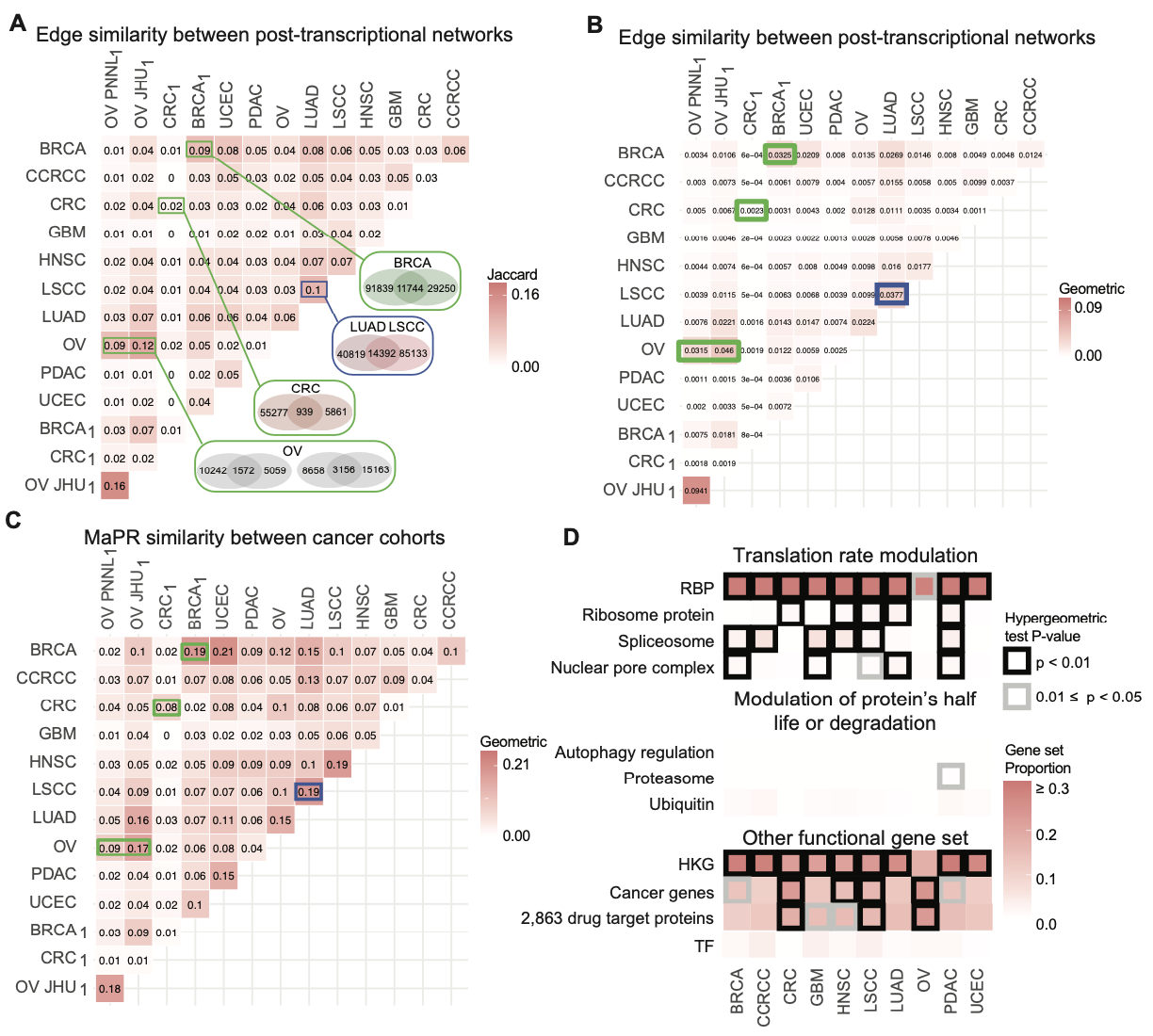

**Figure S2.** (A) The Jaccard index matrix illustrates the similarity of post-transcriptional regulation network across datasets. Same cancer from independent cohort were highlighted in green box while cancers originating from same tissue by blue box, where the number of overlapped post-transcriptional regulation edges is shown. (B-C) The Geometric index matrix delineates the similarity of post-transcriptional regulation network (B) or MaPR (C) across datasets. Geometric index calculated as the multiply of the overlapped proportion between two gene sets. Cancer type from validation dataset were subscript with a ’1’. (D) Overlap of MaPR of each cancer among gene set reported with known functions. Each cell represents the percentage of MaPRs overlapping with these gene sets with known functions. Only significant cells, hypergeometric test P-value < 0.05, were boxed. Cancer genes are the union of cancer hallmark genes (Yize Li et al Cell 023) and genes from the Cancer Gene Census of Cosmic. The 2,863 drug target proteins classified into five tiers were from Savage et al Cell 2024. Abbreviations: RBP: RNA binding protein; HKG: Housekeeping Gene; TF: transcription factor.

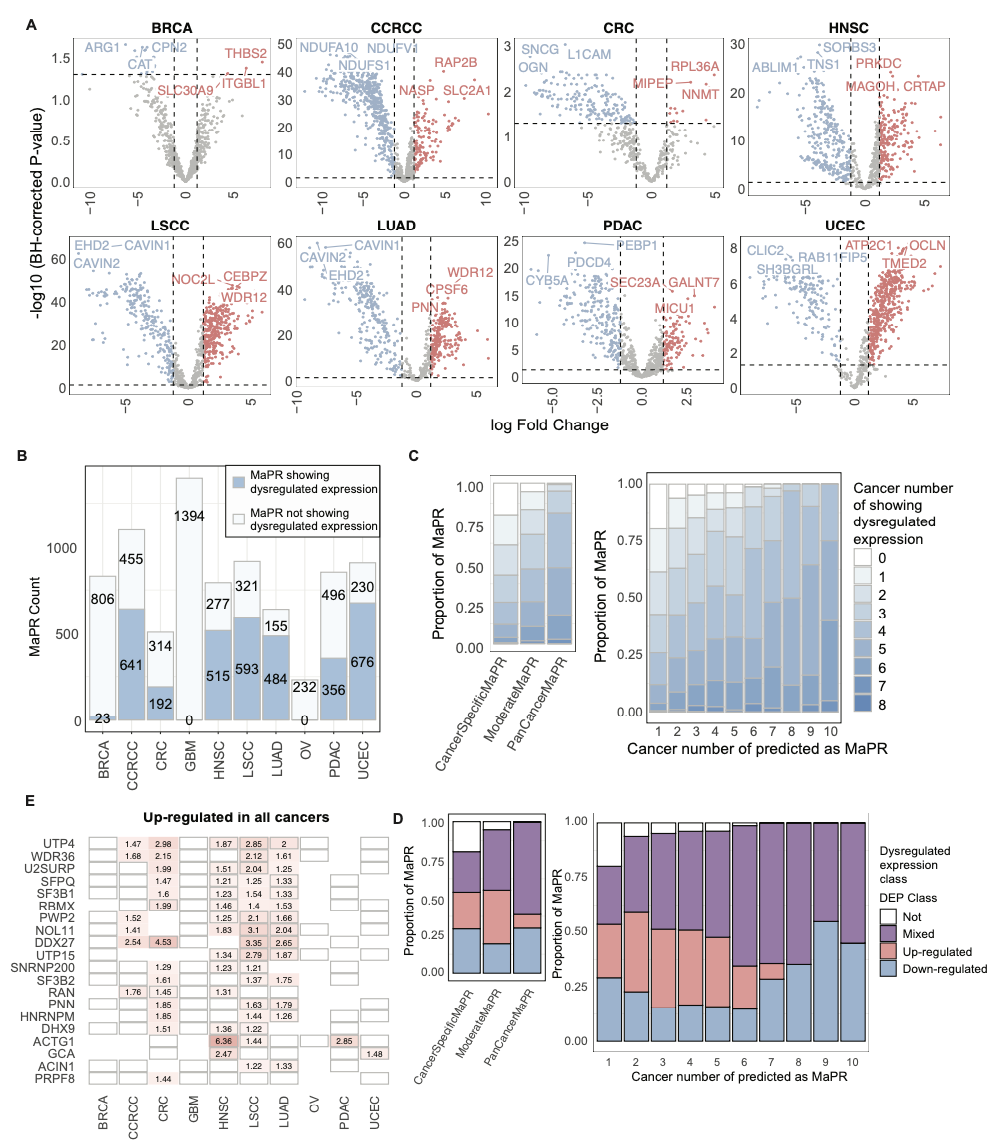

**Figure S3.** MaPR showing dys-regulated protein expression across cancers. GBM and OV are not included due to insufficiency of patient with matched tumor/normal samples to perform differential expression analysis. (A) MaPRs displaying dysregulated protein expression across 8 cancers. X-axis represents log Fold Change in protein expression, with positive indicating up-regulated expression in tumor. Y-axis represent –log10 transform of BH-corrected P-value. Top three MaPR that shows dysregulated protein expression in each cancer were highlighted in red (up-regulated) and blue (down-regulated). (B) The number of MaPRs showing dys-regulated protein expression across cancers. (C) Proportion of MaPRs showing dys-regulated protein expression across three categories (left) or different number level of cancer types (right). Colors represent the number of cancers where a MaPR exhibits significant dys-regulate protein expression. (D) Proportion of MaPRs showing consistent dys-regulated expression direction across three MaPR categories (left) and different number level of cancer types (right). (E) The degree to which pan-cancer consistently up-regulated MaPRs exhibit. The color scale represents the log fold change of expression, with positive values indicating up-regulated expression in tumor. A grey box indicated a protein predicted as MaPR in a specific cancer.

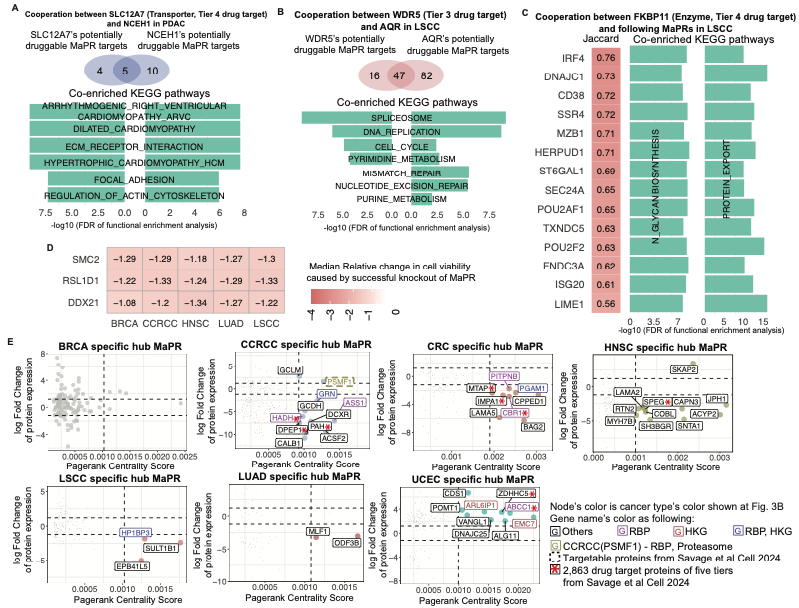

**Figure S4.** (A) Cooperation between SLC12A7, a transporter protein of Tier 4 drug target, and NCEH1 in PDAC. Tier 4 drug target belongs to protein families frequently targeted by small molecules. Venn showed the overlap between two MaPRs’ potentially druggable MaPR targets and their co-enriched KEGG pathways. (B) Cooperation between WDR5, Tier 3 drug target, and AQR in LSCC. Venn showed the overlap between two MaPRs’ potentially druggable MaPR targets and their co-enriched KEGG pathways. Tier 3 drug target were inhibited by drugs considered investigational or experimental. (C) Cooperation between FKBP11, an enzyme of Tier 4 drug target, and other 14 MaPRs in LSCC. Tier 4 drug target belongs to protein families frequently targeted by small molecules. Venn showed the overlap between two MaPRs’ potentially druggable MaPR targets and their co-enriched KEGG pathways. (D) Median relative change in cell viability (Chronos score) caused by successful knockout of three MaPRs among cell lines corresponding to three cancers from DepMap 24Q2. (E) Cancer specific hub MaPRs, defined as MaPR with top 2% Pagerank, that show significant dys-regulated protein expression in specific cancer. Two horizontal dashed lines represent ±1.2 of log Fold Change of protein abundance while one vertical dashed line represents the 0.9 quantile Pagerank score. Name color stands for gene set with known functions (Fig. 2B) while node color represents the cancer type where protein was predicted as MaPR.

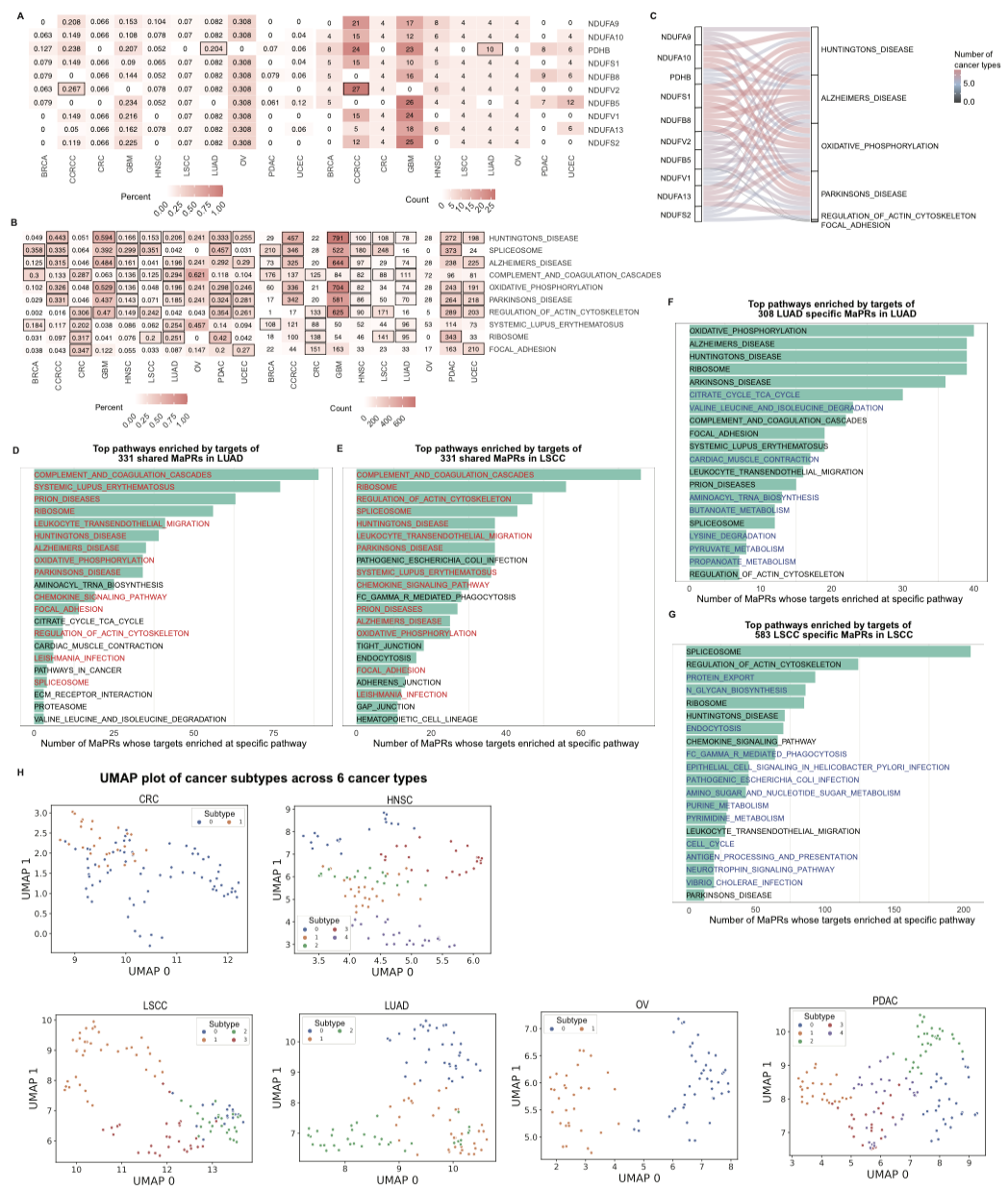

**Figure S5.** (A) In each cancer, one MaPR’s targets enriched at multiple pathways, occupying a proportion of all enriched pathways in that cancer. Shown are top 10 MaPRs with the highest overall proportion across cancers. Boxed cell means that MaPR is also among the top 10 MaPRs in that cancer type. (B) In each cancer, one pathway were enriched by multiple MaPRs’ targets, occupying a proportion of all enriched MaPRs in that cancer. Shown are top 10 pathways with the highest overall proportion across cancers. Boxed cell means that pathway is also among the top 10 pathways in that cancer type. (C) MaPRs of (A) were connected to the pathways of (B). The line width represents the number of cancer types where a MaPR enriched at a pathway. Pathways without enrichment by targets of these MaPRs in any cancer were not shown. (D-G)Top pathways enriched by targets of: (i) 331 shared MaPRs between LSCC and LUAD, (ii) 308 LUAD specific MaPRs, (iii) 583 LSCC specific MaPRs. The shared pathways between (D) and (E) were labelled red. The pathways of (F) not on (E,G) and the pathways of (G) not on (D,F) were labelled blue. X-axis is the number of MaPRs whose targets enriched at specific pathway. (H) Proteomic subtype membership for samples in each cancer type. UMAP was applied to the proteomics data and K-Means clustering was applied to the first 3 UMAP components. Silhouette scores were calculated to find the optimal number of proteomic subtypes for each cancer type. Each sample is colored by proteome subtype membership.

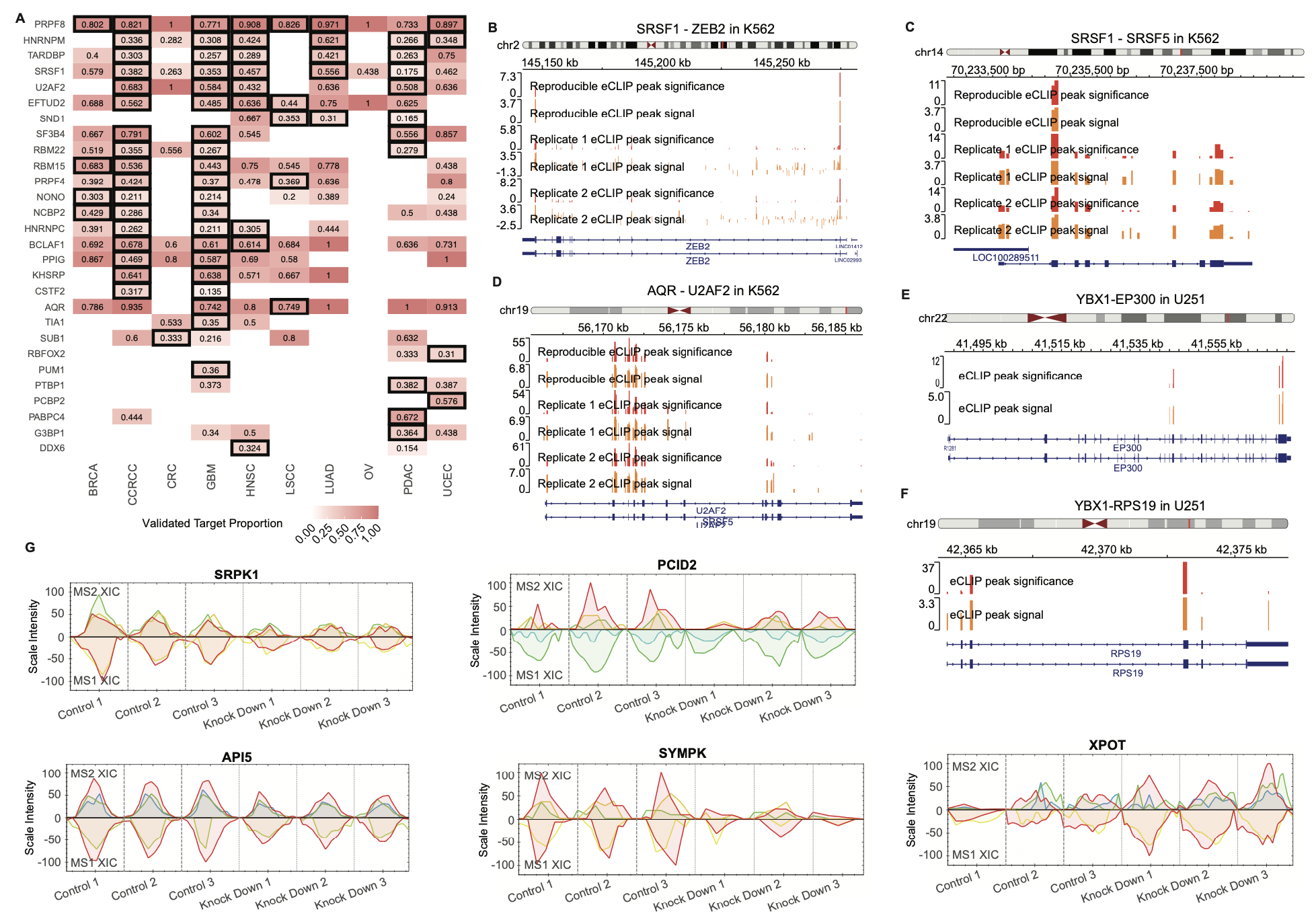

**Figure S6.** (A) Each cell stands for the percentage of predicted targets validated by eCLIP. Only significantly validated cells shown in red and respective validation rate. Only protein predicted as MaPR in a specific cancer is boxed. Only rows with at least one cell > 0.3 were showed here. (B-C) eCLIP RNA binding peaks of SRSF1 on ZEB2 and SRSF5 in K562 cell line. (D) eCLIP RNA binding peaks of AQR on U2AF2 in K562 cell line. (E-F) eCLIP RNA binding peaks of YBX1 on EP300 and RPS19 in glioblastoma cell line of U251. (G) SWATH-MS data upon PRPF8 knockdown in breast epithelial-like cell line of Cal51 shown by XIC graph for five PRPF8 targets: SRPK1, PCID2, API5, SYMPK and XPOT. XIC graph used the extracted ion chromatography to measure protein abundance quantifications in three biologically independent replicate (control-siRNA-treated and PRPF8-depleted). For each XIC graph, the upper panel of peak graphs represent the MS2 level ion traces while the bottom panel of peak graphs represent the MS1 level ion traces.
